# Genetic conflicts between sex chromosomes drive expansion and loss of sperm nuclear basic protein genes in *Drosophila*

**DOI:** 10.1101/2022.06.08.495379

**Authors:** Ching-Ho Chang, Harmit S. Malik

**Author notes:** Correspondence should be addressed to: Ching-Ho Chang, 1100 Fairview Avenue N. A2-205, Seattle, WA 98109.

## Abstract

Many animal species employ short, positively charged proteins, called sperm nuclear basic proteins (SNBPs) or protamines, for tighter packaging of genomes in sperm. SNBP repertoires differ dramatically across animal lineages and signatures of rapid evolution have been reported in mammals. Both sperm competition and meiotic drive between sex chromosomes have been proposed as causes of SNBP innovation. We used a phylogenomic approach to investigate SNBP diversification and its underlying causes in *Drosophila* species. We found unambiguous signatures of positive selection in most SNBP genes except for genes essential for male fertility in *D. melanogaster*. Unexpectedly, evolutionarily young SNBP genes are more likely to encode essential functions for fertility than ancient, conserved SNBP genes like *CG30056*, which we found is dispensable for male fertility despite universal retention in *Drosophila* species. We found 19 independent amplification events involving eight SNBP genes that occurred preferentially on sex chromosomes in 78 *Drosophila* species. Conversely, we found that otherwise-conserved SNBP genes were lost in the *montium* group of *Drosophila* species, coincident with an X-Y chromosomal fusion. Furthermore, SNBP genes that became linked to sex chromosomes via chromosomal fusions are prone to degenerate or relocate back to autosomes. We hypothesize that SNBP genes ancestrally encoded by autosomes suppress meiotic drive, whereas sex-chromosomal SNBP expansions directly participate in meiotic drive. X-Y fusions in the *montium* group render autosomal SNBPs dispensable by making X-versus-Y meiotic drive obsolete or costly. We conclude that SNBP rapid evolution is driven by genetic conflicts between sex chromosomes during spermatogenesis in *Drosophila* species.

## Introduction

Chromatin plays a critical role in organizing genomes and regulating gene expression. Histones are the primary protein components of chromatin in most eukaryotes. Due to their conserved, essential functions, most histone proteins are ancient and subject to strong evolutionary constraints, although there are distinct exceptions among histone variants [1–3]. Many animal species replace most histones with sperm nuclear basic proteins (SNBPs) to pack their genomes during spermatogenesis [4, 5]. This ensures that genomic DNA is packaged even more compactly into tiny sperm heads. Like histones, most SNBPs are small (<200 amino acids) and positively charged. Many SNBPs also contain a high proportion of lysine, arginine, and cysteine residues, which form disulfide bridges to further condense DNA within sperm heads [6, 7]. As a result of their tighter DNA packaging, SNBPs can reduce the size of the sperm nuclei by 20–200 fold compared to histone-enriched nuclei [8]. Based on this tighter packaging, the prevailing hypothesis is that sexual selection for competitive sperm shapes led to the evolutionary origins of SNBP genes in most animal taxa [9, 10].

SNBPs have been most well-studied in mammals [11]. Mammalian SNBPs in mature sperm include protamine 1 (*PRM1*) and protamine 2 (*PRM2*), which are encoded in an autosomal gene cluster that includes Transition Protein 2 (*TNP2*) and protamine 3 (*PRM3*) [12]. Although these four genes share moderate homology, *TPN2* is only expressed during the transition from histones to protamines [13], whereas PRM3 only localizes to the cytoplasm of elongated spermatids [12]. Both *PRM1* and *PRM2* are essential for fertility in humans and mice; their expression levels directly affect sperm quality [11]. Loss of *PRM1* or *PRM2* leads to defects in sperm head morphology and fertility in mice and humans [14, 15] and yet, *PRM2* has undergone pseudogenization in bulls [11]. Thus, even SNBPs essential for male fertility can be subject to evolutionary turnover in some species.

Although SNBPs play a similar genome-packaging role to histones, they differ dramatically from histones in their evolutionary origins and trajectories. Whereas histones have ancient origins, SNBPs arose independently from different ancestral proteins across taxa [7, 16, 17]. For example, many SNBPs arose independently from linker histone H1 gene variants in liverworts and tunicates [18, 19] and from histone H2B gene variants in cnidarians and echinoderms [6, 20, 21]. SNBPs in other animals lack apparent homology to other existing proteins, obscuring their evolutionary origins [21]. In addition to their convergent evolution and turnover, SNBPs also differ dramatically from histones in their evolutionary rates of amino acid divergence. For example, *PRM1* and *PRM2* are among the most rapidly-evolving protein-coding genes encoded in mammalian genomes [22] and evolve under positive selection in many lineages [9, 23]. One of the driving forces leading to the positive selection of SNBPs is changes in their amino acid composition. For example, the arginine content of PRM1 is partially correlated across species with sperm head length, which may reflect the selective pressures of sperm competition [24]. The rapid evolution of *PRM1* and *PRM2* is consistent with sexual selection on sperm heads driving SNBP origins and rapid evolution in mammals [23–25], although this hypothesis has yet to be experimentally tested. However, the evolutionary trajectories of SNBP genes and their underlying causes have not been deeply investigated outside mammals.

*Drosophila* species provide an excellent model to study SNBP function and evolution due to the ease of genetic manipulations and sperm biology characterization, and the availability of high-quality genome sequences from many closely related species. Previous studies have shown that *Drosophila* SNBPs independently arose from proteins encoding high mobility group box (HMG-box) DNA-binding proteins [26]. Five HMG-box SNBP genes have been previously identified in *D. melanogaster: ProtA, ProtB, ddbt, Mst77F,* and *Prtl99C* [27–31]. Each of these five SNBPs is incorporated into nuclei independent of each other, suggesting they play distinct roles in sperm formation [28, 30, 32]. Based on the common HMG-box motifs found in these five SNBPs, ten other male-specific proteins with the same motif have been identified in *D. melanogaster* [26, 30], four of which were later shown to be enriched in sperm nuclei [26]. Some other proteins without any HMB-box are also demonstrated to locate in the sperm nucleus, but it is unclear whether they bind to DNA [33–35].

Recent studies in *Drosophila* have suggested an alternate hypothesis to sperm competition— meiotic drive and its suppression—to explain the rapid diversification and innovation of SNBP-like proteins [36, 37]. Meiotic drivers are selfish elements that can bias their transmission via hijacking meiosis or post-meiosis processes, *e.g.*, killing sperm that do not carry the driver. These genetic drivers exist in widespread lineages, including plants, animals, and fungi [38]. One of the first identified drivers is *Segregation Distorter* in *D. melanogaster*, whose drive strength can be enhanced by the knockdown of *ProtA/ProtB* genes [39]. Thus, *ProtA/ProtB* serve as suppressors of meiotic drive through an unknown mechanism. The second piece of evidence emerged from studies of *Distorter on the X (Dox),* an X-chromosomal driver in *D. simulans* [40]. *Dox* emerged via the stepwise acquisition of multiple gene segments, mostly from *ProtA/ProtB*. *Dox* produces chromosome condensation defects in Y chromosome-containing sperm during spermatogenesis, ultimately leading to X-chromosomal bias among functional sperm and sex-ratio bias in resulting progeny [40–42]. In *D. simulans* and sister species, *Dox*-like genes have amplified and diversified on the X chromosome in an escalating battle between X and Y chromosomes for transmission through the male germline [36, 37]. Thus, genetic conflicts between sex chromosomes and their suppression of those conflicts provide an alternate explanation for the recurrent diversification of SNBP genes in *Drosophila* species.

Here, we systemically explored the evolution of SNBP genes via a detailed phylogenomic analysis across *Drosophila* species. We found that SNBP genes are rapidly evolving and most of them are under positive selection in *Drosophila*, consistent with the observation in mammals. Thus, the rapid sequence changes of SNBP genes are common to many animal taxa. Interestingly, we find an inverse relationship between age and essentiality; young SNBPs are essential for male fertility in *D. melanogaster*, whereas ancient, conserved SNBPs are not. Moreover, SNBP genes essential for male fertility in *D. melanogaster* are frequently lost in other *Drosophila* species. Unexpectedly, we found 19 independent amplification events from eight different SNBP genes on either X or Y chromosomes in *Drosophila* species. Conversely, species with reduced conflicts between sex chromosomes due to chromosomal fusions do not undergo SNBP amplification, and instead, frequently lose SNBP genes. Thus, we conclude that rapid diversification of SNBP genes might be largely driven by genetic conflicts between sex chromosomes in *Drosophila*.

## Results

### SNBP genes in *Drosophila* species

To study SNBP evolution in *Drosophila* species, we performed a detailed survey of all testis-specific genes encoding HMG boxes in *D. melanogaster.* Our survey did not reveal any additional genes beyond the 15 previously identified candidate SNBP genes, which function at different stages of spermatogenesis (Table 1). For example, *CG14835, ProtA, ProtB, Mst77F, Prtl99C,* and *ddbt* all encode SNBP proteins present in the mature sperm head [29–31]. In contrast, *Tpl94D, tHMG-1, tHMG-2, and CG30356* encode transition SNBP proteins during the transition between histone and protamines but are not retained in mature sperm [26, 43]. The five remaining SNBP genes (*Mst33A, CG30056, CG31010, CG34269, and CG42355*) remain cytologically uncharacterized [30]. SNBP proteins in *D. melanogaster* tend to be short (<200 a.a.) and mostly have high Isoelectric points (>10), consistent with their basic charge and potential function in tight packaging of DNA (Table 1). A closer examination revealed that eleven SNBP genes encode a single HMG box, whereas four genes (*Tpl94D, Prtl99C, Mst33A* and *CG42355*) encode two HMG boxes (Table 1 and Fig S1).

**Table 1.**
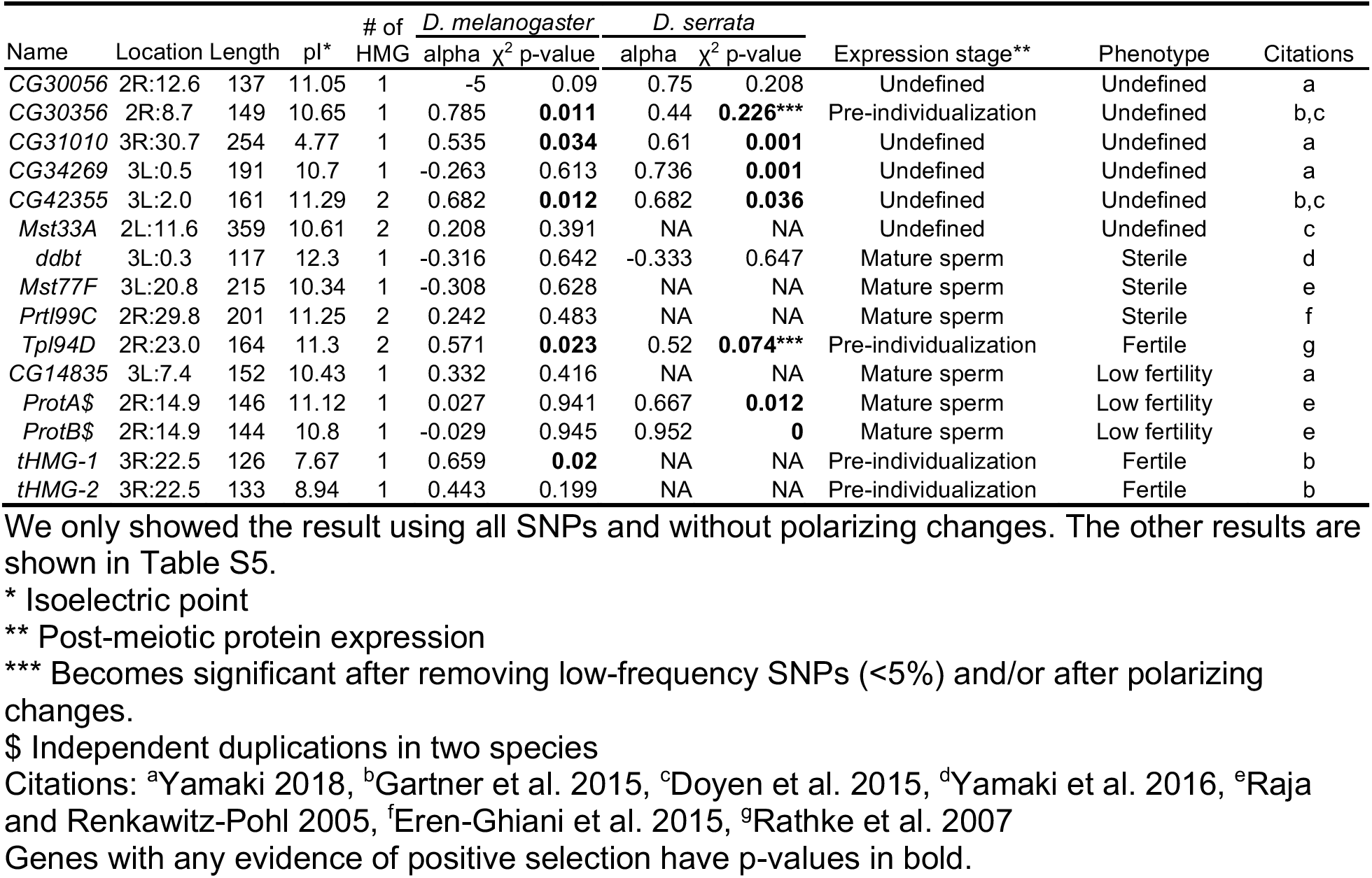
McDonald–Kreitman tests for positive selection on SNBP genes in two Drosophila species.

To investigate the retention of SNBP genes across *Drosophila* species, we expanded our analysis to homologs of *D. melanogaster* SNBP genes found in published genome assemblies from 15 *Drosophila* species with NCBI gene annotation. We also included *Scaptodrosophila lebanonensis* as an outgroup species. Our phylogenomic analyses revealed that two SNBP genes (*tHMG* and *Prot*) underwent very recent gene duplications, specifically in *D. melanogaster*. Both are present as closely related paralogs (*tHMG-1* and *tHMG-2, ProtA* and *ProtB*) in *D. melanogaster* but only in one copy in closely related species (Fig 1) [27, 29]. Five SNBP genes are found only in the *Sophophora* subgenus: *CG42355, Mst33A, Mst77F, Prtl99C,* and *Tpl94D* (Fig 1), and are therefore, less than 40 million years old. At the other extreme, we found orthologs of eight *D. melanogaster* SNBP genes (*CG14835, CG30056, CG30356, CG31010, CG34529, ddbt, tHMG* and *Prot*) in the outgroup species, *S. lebanonensis*. Thus, these eight SNBP genes are at least 50 million years old [44]. Our analysis focuses on SNBP genes present in *D. melanogaster*, but other *Drosophila* species may have additional, unrelated SNBP genes.

**Figure 1.**
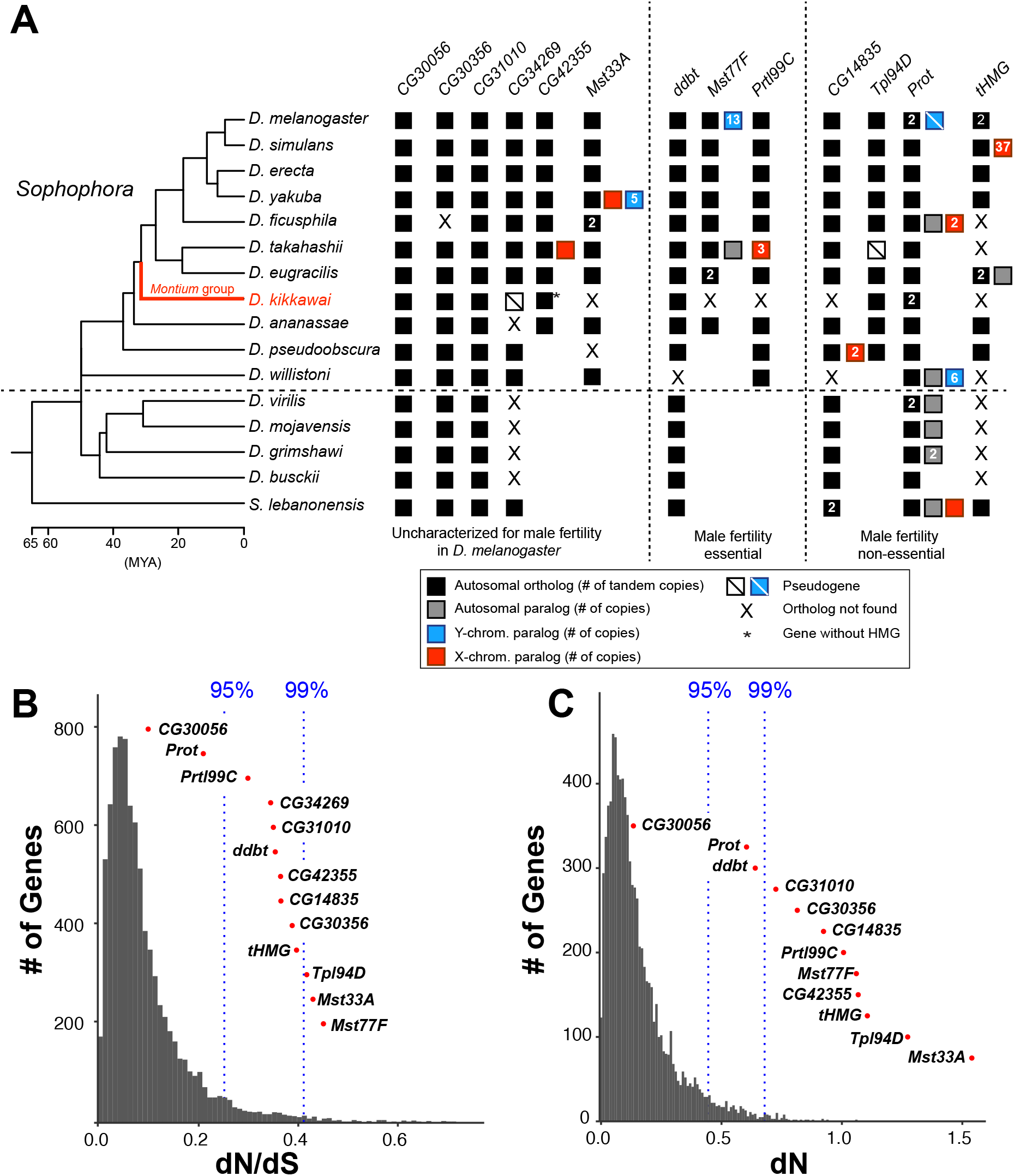
Origins and evolution of *Drosophila* SNBP genes. **(A)** Phylogenomic analysis of 13-15 SNBP genes from *D. melanogaster* organized into three groups (dotted lines)-either required for male fertility, not required for male fertility, or untested in previous analyses. We identified homologs of these genes in 14 other *Drosophila* species and an outgroup species, *S. lebanonensis*, whose phylogenetic relationships and divergence times are indicated on the left [88]. Genes retained in autosomal syntenic locations are indicated by black squares, whereas paralogs located in non-syntenic autosomal locations, or X-chromosomes, or Y-chromosomes are indicated in gray, blue and red squares, respectively. Numbers within the squares show the copy number, if >1 of different genes, *e.g., D. melanogaster* has two paralogs of both *Prot* and *tHMG* genes. An empty square with a line across it indicates that only a pseudogene can be found, whereas an ‘X’ indicates that no ortholog is found, even though one is expected based on the phylogenomic inference of SNBP age. Based on this analysis, we infer that eight SNBP genes are at least 50 million years old, although only two genes are strictly retained in all 16 species. Indeed, none of the SNBP genes required for male fertility in *D. melanogaster* are strictly conserved in other *Drosophila* species, either arising more recently (*Mst77F, Prtl99C*) or ancient but lost in at least one species (*ddbt*). We also marked the *montium* group species, *D. kikkawai*, in red, losing eight SNBP genes. **(B)(C)** We compared dN/dS **(B)** or dN **(C)** values for all orthologous SNBP genes (red dots) in *D. melanogaster* compared to a histogram of the same values for the genome-wide distribution (gray bars) obtained from an analysis using six species by the 12 Drosophila genomes project [72]. Our analyses reveal that most SNBP genes are at or beyond the 95^th^ or 99^th^ percentile for dN/dS or dN values (blue dashed lines). The values of *CG34269* are calculated using only five species because it is lost in one of the surveyed species, *D. ananassae*, and therefore we do not show its dN, as it is not comparable to other genes.

Our inability to detect homologs beyond the reported species does not appear to result from their rapid sequence evolution. Indeed, abSENSE analyses [45] support the finding that *Prtl99C, Mst77F, Mst33A, Tpl94* and *CG42355* were recently acquired in *Sophophora* within 40 MYA. For example, the probability of a true homolog being undetected for *Prtl99C* and *Mst77F* is 0.07 and 0.18 (using E-value=1), respectively (**Table S1**, Methods). Although abSENSE analyses can rule out the absence of true homologs, they cannot rule out the less parsimonious possibility that SNBP genes are older but were lost multiple times in non-*Sophophora* species.

In addition to orthologs of these SNBP genes found in shared syntenic locations, we also found sex chromosome-linked paralogs of SNBP genes in several species. The most dramatic example is the presence of 34 copies of *tHMG* paralogs in the poorly-assembled X chromosomal region of *D. simulans* (Fig 1). These are discussed in more detail later in this study.

We confirmed that *Drosophila* SNBP gene expression is primarily male-limited across species using publicly available RNA-seq data; their expression is particularly enriched in testes (Fig S2; Table S2). The only exception is a *CG42355* paralog in *D. takahashii* that also has weak expression in females (∼9 TPM; Fig S2B; Table S2). We observed a medium to high correlation (Spearman’s rho= 0.142–0.753; Fig S3) for the expression of SNBP genes between species. Like in *D. melanogaster*, most *Drosophila* SNBP proteins are small, possess at least one HMG box domain and have high isoelectric points, suggesting these features are crucial for their function (Table S3).

### Rapid evolution and positive selection of *Drosophila* SNBP genes

Based on the precedent of rapid evolving protamines in mammals, we next investigated whether *Drosophila* SNBP genes also evolve rapidly. We calculated protein evolution rates (nonsynonymous substitution rates over synonymous substitution rates, dN/dS) for 13 of 15 *D. melanogaster* SNBP genes for six species in the *melanogaster* group (Table S4). We excluded two SNBP genes, *ProtB* and *tHMG-2,* since these duplicates are not found outside *D. melanogaster*. We found that 11 of 13 SNBP genes (except *CG30056* and *ProtA*) evolve faster (higher dN/dS) than 95% of protein-coding genes across the genome (Fig 1B). These high protein evolution rates are due to high dN instead of low dS (Fig 1C), suggesting that SNBPs evolve under extensive positive selection or reduced functional constraints.

We used McDonald–Kreitman tests to test the possibility of recent positive selection in the *D. melanogaster* lineage, taking advantage of many sequenced strains from this species [43, 44, 49-56]. The McDonald-Kreitman tests compare the ratio of non-synonymous to synonymous substitutions fixed during inter-species divergence to the ratio of these polymorphisms segregating within species. If there is an excess of non-synonymous changes fixed during inter-species divergence, this results from positive selection. Indeed, our tests reveal that five SNBP genes in *D. melanogaster* have evolved under positive selection during its divergence from *D. simulans* (Table 1). By polarizing the test using the inferred ancestral sequences of *tHMG-1* and *tHMG-2,* we find that *tHMG-1*, but not its paralog, *tHMG-2,* evolved under positive selection in *D. melanogaster* (Table S5).

We also took advantage of genome sequences from 110 *D. serrata* strains to carry out McDonald-Kreitman tests of SNBP genes from *D. serrata* compared to its sister species, *D. bunnanda* in the *montium* group [46, 47]. Among the seven SNBP genes shared between *D. melanogaster* and *D. serrata*, four genes (*CG30356, CG31010, CG42355, Tpl94D*) evolved under positive selection in both *D. melanogaster* and *D. serrata*, whereas two genes (*CG30056, ddbt*) do not show a signature of positive selection in either species. Three additional genes (*CG34269,* and two *ProtA/B* duplicates) evolved under positive selection only in *D. serrata* (*ProtA/B* underwent independent gene duplications in *D. melanogaster* and *D. serrata*).

Finally, we used maximum-likelihood analyses using the site model of the codeml program in PAML package [48, 49] to investigate whether any residues in the SNBP genes had evolved under recurrent positive selection. We limited our analyses to 17 species of the *melanogaster* group, to avoid saturation of synonymous substitutions. Although we recapitulated a previously published positive selection result using *ddbt* genes from just five *Drosophila* species [31], analyses using 17 *melanogaster* group species did not find the site-specific positive selection in any SNBP gene (Table S6). The discrepancy between the McDonald-Kreitman tests and the PAML results indicate that although many SNBP genes evolve under positive selection, either the positive selection of SNBP genes or the exact residues evolving under recurrent positive selection varies throughout *Drosophila* evolution.

### What determines SNBP essentiality for male fertility?

Nine of fifteen SNBP genes have been previously characterized for their roles in male fertility based on gene knockout or knockdown experiments in *D. melanogaster* (Table 1). These genes show differences in their importance for male fertility in *D. melanogaster*. Three genes (*Mst77F*, *Prtl99C* and *ddbt*) are essential for male fertility while knockdown or knockout of two other SNBP genes (*ProtA,* and *ProtB*) leads to significant reduction in male fertility [27–30, 32]. However, knockdown of four genes (*CG14835, Tpl94D, tHMG-1*, and *tHMG-2*) does not appear to impair male fertility at all. No information is currently available for the remaining six SNBP genes. We also found nearly strict retention of all SNBP genes in all sequenced strains of *D. melanogaster,* no matter whether they are essential for male fertility in laboratory experiments or not (Table S7 and Supplementary text).

There are a few distinguishing characteristics common to SNBP genes required for male fertility. Neither the number of HMG domains nor expression levels of SNBP are associated with essentiality. Instead, proteins that are essential (*Mst77F*, *Prtl99C* and *ddbt*) or important (*ProtA* and *ProtB*) for male fertility are more likely to be present in the mature sperm head, whereas transition SNBPs (*Tpl94D, tHMG-1*, and *tHMG-2*) are more likely to be dispensable, potentially due to functional redundancy. Moreover, SNBP genes important for male fertility in *D. melanogaster* show no signature of positive selection according to McDonald-Kreitman tests. In contrast, two of the three identified transition SNBP genes evolve under positive selection (*Tpl94D* and *tHMG-1*). This suggests that genes with redundant function or less critical for male fertility are more likely to evolve under positive selection, although we note that several SNBP genes remain functionally uncharacterized or have not been tested exclusively (Table 1).

How does SNBP essentiality in *D. melanogaster* correlate with age and retention across *Drosophila* species evolution? We find that two essential SNBP genes (*Prtl99C* and *Mst77F*) are evolutionarily young *i.e.,* they arose relatively recently in *Drosophila* evolution. Moreover, both genes have been lost at least once since their birth, in *D. kikkawai*. The third essential SNBP gene, *ddbt*, arose before the separation of *Drosophila* and *S. lebanonensis*, but it has also been lost at least once (in *D. willistoni*) among the 15 species analyzed [31] (Fig 1A). Based on these findings, we infer that not only are these three essential SNBP genes subject to evolutionary turnover, but they also gain or lose essential function across *Drosophila* evolution. Our findings are similar to recent studies that show high evolutionary lability of many genes involved in essential heterochromatin or centromere function in *Drosophila* [50, 51].

### *CG30056* is universally retained in *Drosophila* but dispensable for male fertility in *D. melanogaster*

Given the high evolutionary turnover of SNBP genes in our sampling of 15 *Drosophila* species, we investigated whether any SNBP genes are universally retained across all *Drosophila* species. For this purpose, we expanded our phylogenomic analysis of SNBP evolution to a recently published dataset of 78 highly contiguous and complete *Drosophila* genomes [52], using tBlastn and reciprocal blastx searches [53]. Based on this analysis, we find only two SNBP genes have been strictly retained across all *Drosophila* species — *Prot* and *CG30056* (Fig S4), which also have the lowest dN/dS in SNBP genes (Fig 1B). Previous studies have shown that loss of *ProtA* and *ProtB* paralogs reduce male fertility in *D. melanogaster* [27].

Despite being one of the most well-retained and highly conserved SNBP genes in *Drosophila* species (Fig 1B), *CG30056* remains functionally uncharacterized for its role in male fertility. To study its contribution to male fertility, we generated a complete deletion knockout of *CG30056* gene using CRISPR/Cas9 (Fig 2A). *CG30056* is located in an intron of *frazzled*, a gene essential for development and morphogenesis (Fig 2A). We co-injected a construct encoding two gRNAs designed to target sequences immediately flanking *CG30056* together with a repair construct containing 3xP3-DsRed (a visible eye marker). We successfully obtained transgenic flies encoding DsRed and confirmed the deletion of *CG30056* using PCR (Fig 2B). Homozygous knockout flies were viable, assuring that removal of *CG30056* did not disrupt the essential *frazzled* gene.

**Figure 2.**
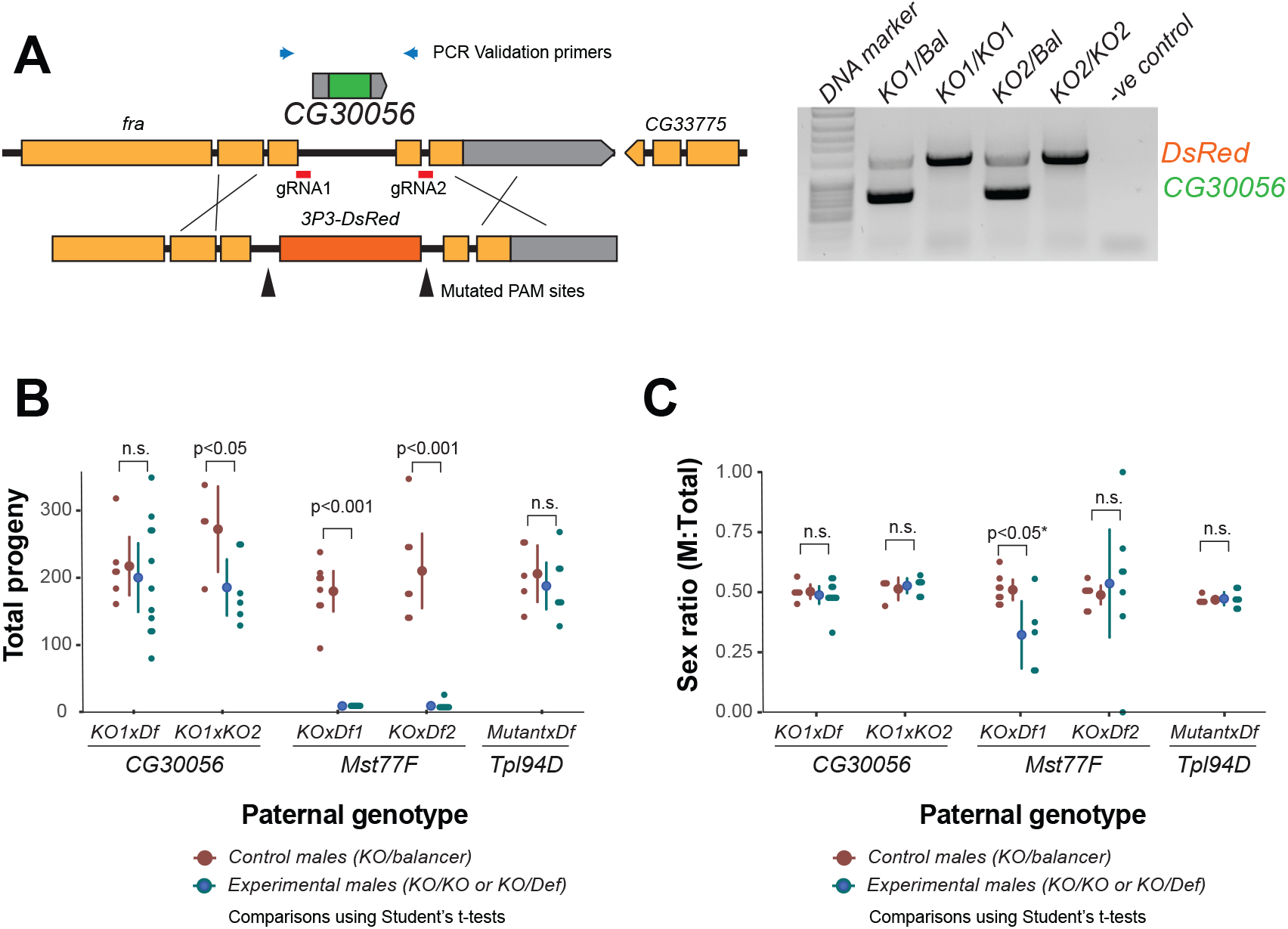
The strictly retained, highly conserved SNBP gene, *CG30056*, is not required for male fertility in *D. melanogaster*. **(A)** The SNBP gene, *CG30056*, is encoded co-directionally in the essential *frazzled* gene intron. Using guide RNAs designed to match sites flanking CG30056, and a healing construct encoding eye-specific DsRed, we created a knockout allele replacing CG30056 with DsRed. The knockout was verified using PCR and primers flanking the CG30056 locus (right). Note that balancer lines encode a wildtype copy of *CG30056*. **(B)** We performed fertility assays comparing *CG30056* homozygous knockout flies with heterozygous controls. We also assayed fertility of knockout strains for the fertility-essential *Mst77F* gene, and the fertility nonessential *Tpl94D* gene, together. We also documented the sex-ratios of the resulting progeny in (C). Consistent with previous findings, we found that Mst77F knockout males are essentially sterile and *Tpl94D* knockout males were indistinguishable from their heterozygous controls. We found either no or weak evidence of fertility impairments in two different crosses with homozygous *CG30056* knockout males. **(C)** We observe no significant evidence of sex-ratio distortion that would suggest an X-versus-Y meiotic drive in progeny resulting from either *CG30056, Mst77F,* or *Tpl94D* knockout males. *Although there is suggestive evidence of sex-ratio distortion in progeny of one of the *Mst77F* genotypes, we note that this is most likely due to stochastic effects of having very few resulting progeny.

Next, we tested the dependence of male fertility on *CG30056* by crossing two different *CG30056*-*KO* founder lines to each other or to a *D. melanogaster* strain with a large deletion spanning *CG30056* (*Df(2R)BSC880*) to obtain *CG30056-KO* homozygous males. As controls, we used *CG30056/SM6a* or *CG30056/CyO* heterozygous males (*SM6a* and *CyO* are balancer chromosomes with an intact copy of *CG30056)*. We compared fertility of these males by mating them with two wildtype females at room temperature and counting their adult offspring. Although *CG30056* is the most conserved SNBP we surveyed, we found no clear difference in offspring number between heterozygous controls and homozygous knockout males (Fig 2B). We also detected no evidence for sex-ratio distortion in our crosses (Fig 2C). As controls, we conducted parallel experiments with mutants of *Mst77F,* previously shown to be essential for male fertility, and mutants of *Tpl94D,* previously shown to be nonessential. Our results recapitulated previous studies, confirming the essentiality of *Mst77F* and the dispensability of *Tpl94D* (Fig 2B) [26, 28]. Thus, despite its strict retention for more than 60 million years of *Drosophila* evolution, *CG30056* is not required for male fertility in *D. melanogaster,* at least under standard laboratory conditions. However, it is possible that *CG30056* might play another function that we did not assay for, such as in sperm storage or precedence.

Thus, two of three SNBP genes (*Mst77F* and *Prtl99C*) essential for male fertility are evolutionarily young and poorly retained, whereas the most sequence-conserved and well-retained SNBP genes (*CG30056*) is not essential. Our work suggests that young SNBP genes are more likely to encode essential, non-redundant male fertility functions than ancient, well-retained ones.

### Recurrent amplification of a subset of SNBP genes on sex chromosomes

Our analysis of SNBP genes across a limited set of *Drosophila* species had already revealed significant evidence of evolutionary turnover (Fig 1). We further analyzed evolutionary turnovers of SNBP genes in 78 *Drosophila* species and two other *Drosophilidae* species, most of which lack detailed gene annotation [52]. We inferred gains and losses of SNBP genes in these species, which we represent on a circular phylogram of all 80 species. To assign the chromosomal location of SNBP genes, we estimated coverage of publicly available Illumina and Nanopore reads (represented in Fig 3 and Table S8) of male or mixed-sex flies from various *Drosophila* species. We also assigned location to specific Muller elements location based on 3285 BUSCO (Benchmarking Universal Single-Copy Orthologs) genes located on the contigs [54]. For species where sequencing reads are available from male and female flies separately, we could readily assign the male-specific regions to the Y chromosomes. However, we could only ascribe a sex-chromosomal linked location for species if no data was available from either BUSCO genes or females (only males and mixed-sex flies).

**Figure 3.**
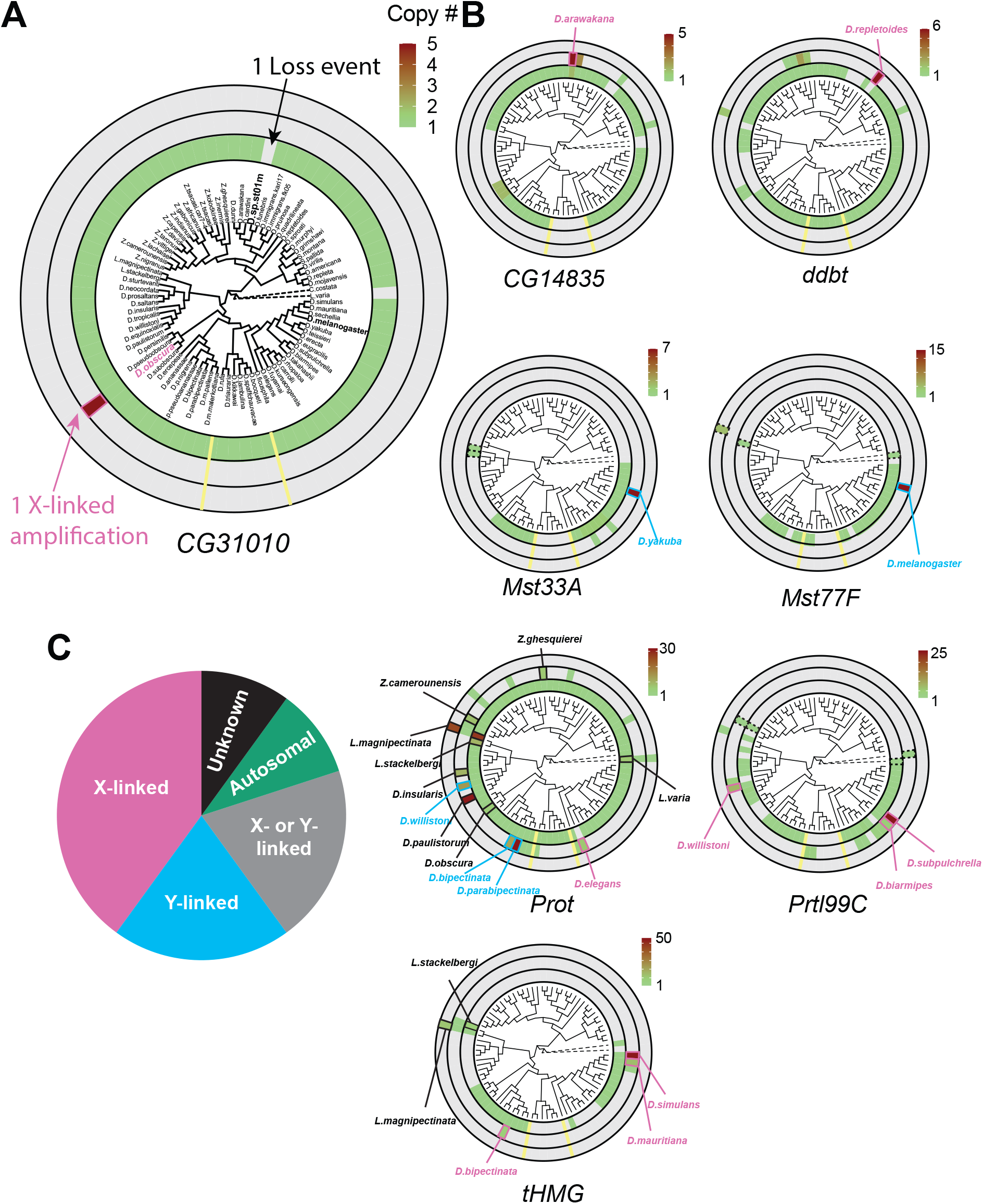
Recurrent amplifications of *Drosophila* SNBP genes are biased for sex-chromosomal linkage. **(A)** Using reciprocal BLAST (see Methods), we searched for homologs of each *D. melanogaster* SNBP gene in 78 distinct *Drosophila* species and two outgroup species (shown in dot lines). We depict our findings using the circular representation indicated in (A) for SNBP gene *CG31010*. The innermost circle is a circular phylogeny of the species [52]. The next circle ring indicates autosomal copies, with colors to indicate copy number (scale bar, top left; note that scales are different for each gene). Thus, *CG31010* is present in one autosomal copy in all but one *Drosophila* species (gray bar). The third circle indicates sex-chromosomal copies. Red and blue frames in the middle ring indicate X- or Y-linkage if that can be reliably assigned. Dotted frames indicate copies that might not be real orthologs based on phylogeny, whereas solid frames indicate five or more copies. For example, *CG31010* is present in 5 copies on the X-chromosome of *D. obscura*. The outermost circle shows copies with ambiguous chromosomal location: there are no such copies for *CG31010*. **(B)** Using the same representation scheme, we indicate gene retention and amplification for six other SNBP genes for which we find robust evidence of amplification, from a copy number of 5 (*CG14835*) to nearly 50 (*tHMG*). We also marked the *montium* group species which lost many SNBP genes with yellow lines. We note that assemblies of *Lordiphosa* species have lower quality, and the data need to be interpreted carefully. **(C)** SNBP gene amplifications (5 or more copies) are heavily biased for sex chromosomal linkage. Given the relative size of sex chromosomes and autosomes, this pattern is highly non-random (test of proportions P=2.3e-5).

All surveyed SNBP genes are located on autosomes in *D. melanogaster* and most species due to rare inter-chromosomal movement of genes in *Drosophila*. Losses or duplications of the autosomal (majorly original) locus are represented in the innermost circle, gains on the sex chromosomes are represented in the next (middle) concentric circle, and gains with ambiguous chromosomal location are represented in the outermost concentric circle (Fig 3A and Fig S4). We use the *CG31010* gene to illustrate this representation. *CG31010* has been retained in all *Drosophila* species except one, which is shown by a gray bar in the innermost concentric circle. In addition, *CG31010* amplified to a total of five X-linked copies in *D. obscura,* represented by a dark red bar in the middle concentric circle (Fig 3A).

Our expanded survey reinforced our initial findings (Fig 1A) that multiple SNBP genes are subject to lineage-specific amplifications (more than five copies in one species). We found that eight of 13 SNBP genes investigated (*CG14835, ddbt, Mst33A, Mst77F, Prtl99C, Prot, tHMG,* and *CG31010)* underwent amplification in at least one species (Fig 3 and Table 2). In total, we found that SNBP genes have undergone 20 independent amplification events, including one event in the outgroup species, *Leucophenga varia* (Fig 3A–B). Most SNBP amplifications are evolutionarily young (< 10 million years old; Fig S5), and 15 of them are specific to a single surveyed species (Fig 3A–B). Many of these amplified copies are arranged in tandem arrays, whose sequences are hard to assemble; our reported numbers are likely an underestimate. Moreover, some amplified SNBP genes, *e.g.*, *Dox*-related genes derived from *ProtA/B* in *D. simulans* [36, 37], are too diverged from the parental genes for unambiguous assignment, and are missing in our survey. Analysis of available RNA-seq datasets revealed that all SNBP paralogs have male-specific, and mostly testis-specific, expression (Fig S2B).

**Table 2.**
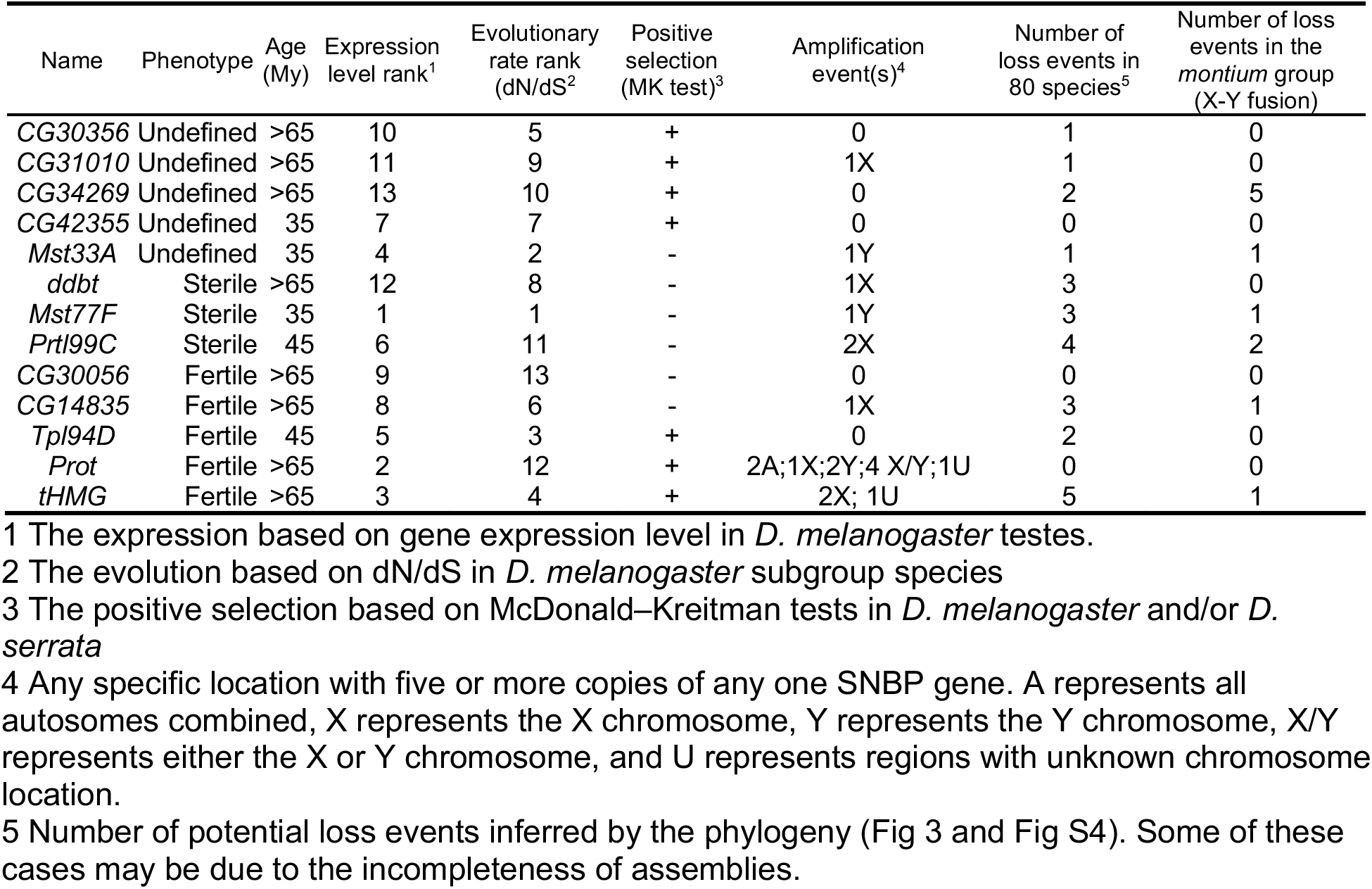
Summary of evolution events of *Drosophila* SNBP genes.

We found that eight amplifications are X-linked, four are Y-linked, whereas for four amplifications, we cannot distinguish between X- or Y-linkage (Fig 3 and Table S9). We infer that 80% (16/20, Fig 3B) of amplifications occurred on sex chromosomes. This high fraction is significantly higher than the null expectation if SNBP amplifications occurred randomly on sex chromosomes or autosomes (∼33% should be on sex chromosomes, given each chromosome arm, except the dot chromosome, has a similar size, Test of Proportions P = 2.3e-5).

To better understand the evolutionary origins and potential function of SNBP gene amplifications, we investigated the amplification of *tHMG* in *D. simulans* and sister species (Fig 4; Supplementary text). We inferred that *tHMG* copies on the heterochromatic X chromosome (*tHMG-hetX)* experienced two duplications before the amplification: first to the euchromatic X region, proximal to *CG12691,* and a subsequent duplication spanning X-linked *tHMG* and part of *CG12691* to pericentromeric X (Fig 4A). We also found 19 and 4 nonsynonymous changes within 81 amino acid residues in *tHMG-hetX* and parental *tHMG* copies (*tHMG-Anc*), respectively (Fig 4B). Our branch test using PAML further showed that both branches have significantly higher protein evolution rates (Fig 4C; ω=1.6, LRT test, P= 0.007; Table S10). although the branch-site test did not reveal the clear evidence of positive selection (LRT test, P= 0.23; Table S10).

**Figure 4.**
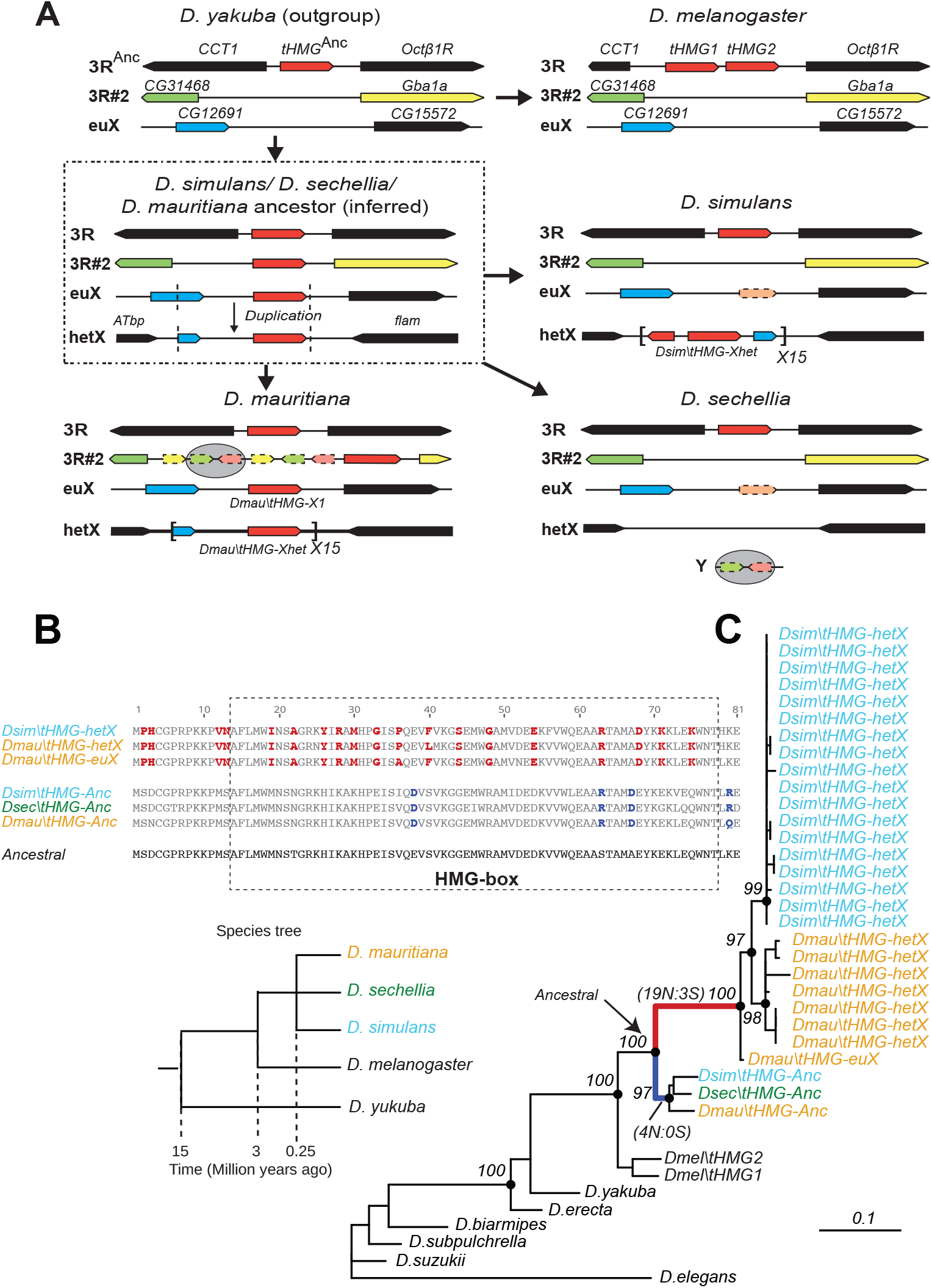
Tracing the duplication and amplification of *tHMG* genes in *D. simulans* and close relatives. **(A)** Using a combination of genome assembly and phylogenetic analyses, we traced the evolutionary origins and steps that led to the massive amplification of *tHMG* genes on the *D. simulans* X chromosome. The first step in this process was the duplication of the ancestral *tHMG* gene (flanked by *CCT1* and *Octb1R*) on the 3R chromosomal arm to a new location on 3R (*tHMG-3R#2* now flanked by *CG31468* and *Gba1a*) and to a location on the X chromosome euchromatin, where *tHMG-euX* is flanked by *CG12691* and *CG15572*. We infer that this *CG12691-tHMG-euX* then duplicated to another locus in X-heterochromatin, between *Atbp* and the *flamenco* locus, and further amplified. These resulting copies experienced different fates in *D. simulans* and its sibling species. For example, in *D. sechellia*, *tHMG-3R#2, tHMG-euX*, and *tHMG-hetX* were all lost but a degenerated copy of *tHMG-3R#2* and flanking genes can be found on its Y chromosome. In contrast, in *D. mauritiana*, *tHMG-3R#2* pseudogenized on 3R, *tHMG-euX* was retained while *tHMG-hetX* underwent an amplification to a copy number of 15 tandemly arrayed genes in the X heterochromatin. Finally, in *D. simulans*, *tHMG-3R#2* was completely lost, *tHMG-euX* was pseudogenized, and *tHMG-hetX* amplified to a copy number of 15 on the X heterochromatin. We note that the amplification unit sizes are different between *D. simulans* and *D. mauritiana*, and between different *D. simulans* strains, suggesting these were independent amplifications. **(B)** The alignment shows the divergence between different *tHMG* copies in the *D. simulans* clade and *D. melanogaster*. We marked 19 and 4 amino acid changes shared among X-linked *tHMG* copies (*tHMG-euX* and *tHMG-hetX*) and parental *tHMG* copies (*tHMG-Anc*), respectively. Most changes are located in the DNA-binding HMG box. **(C)** Phylogenetic analyses of the various *tHMG* genes confirm the chronology of events outlined in (A) and find strong evidence of concerted evolution among the amplified *tHMG-hetX* copies on *D. mauritiana* and *D. simulans*. For comparison, we showed the species tree on the right, and the phylogeny of three *D. simulans* clade species is not solved due to lineage sorting and gene flow. To simplify the analysis, we only used sequences with annotation in NCBI.

### Short life of sex chromosome-linked SNBP genes

Previous studies have shown that the *Dox* meiotic driver arose from an SNBP partial gene amplification (*ProtA/B*) in *D. simulans* [36, 37]. We hypothesize that the *Drosophila* sex-chromosomal SNBP amplifications we have found might similarly be involved in genetic conflicts between sex chromosomes across the *Drosophila* genus, via X-versus-Y meiotic drive. In contrast, all ancestral single-copy SNBP genes that gave rise to these amplifications are encoded on autosomal loci and thus are more likely to encode suppressors of meiotic drive, as is the case for the *ProtA/B* genes against *Segregation Distorter* [39]. If our hypothesis for this duality of SNBP gene functions is correct, we would further predict that ancestrally autosomal SNBP genes that became linked to sex chromosomes would preferentially amplify to become meiotic drivers or be lost due to loss of ancestral functional requirements. To test this hypothesis, we surveyed SNBP genes that became linked to sex chromosomes via chromosome fusions. We found three SNBP genes, *CG14835*, *CG34269* and *ddbt*, which are widely retained in *Drosophila* species on the Muller D element, which is ancestrally autosomal. Both *CG14835* and *ddbt* genes independently became X-linked in the *D. willistoni*, *D. pseudoobscura,* and *D. repletoides* clades, while *CG34269* became X-linked in the two former clades. Consistent with our hypothesis, some newly X-linked SNBP genes either degenerated (2 instances out of 8) or translocated back to autosomes (1 instance out of 8). In five cases, SNBP genes were still retained on sex chromosomes (Fig 5 and Table S11). Among these five cases, we observed one amplification event– *ddbt* amplified to six copies in *D. repletoides* (Fig 3B), consistent with it now acting as a meiotic driver. In contrast, we found no instances of pseudogenization or subsequent translocation to the X chromosome of SNBP genes that are still preserved on their original autosomal locations or involved in chromosome fusions between autosomes (0/16). This difference is highly significant (Fig 5 and Table S11; 3:5 versus 0:16, Fisher’s exact test, P=0.03).

**Figure 5.**
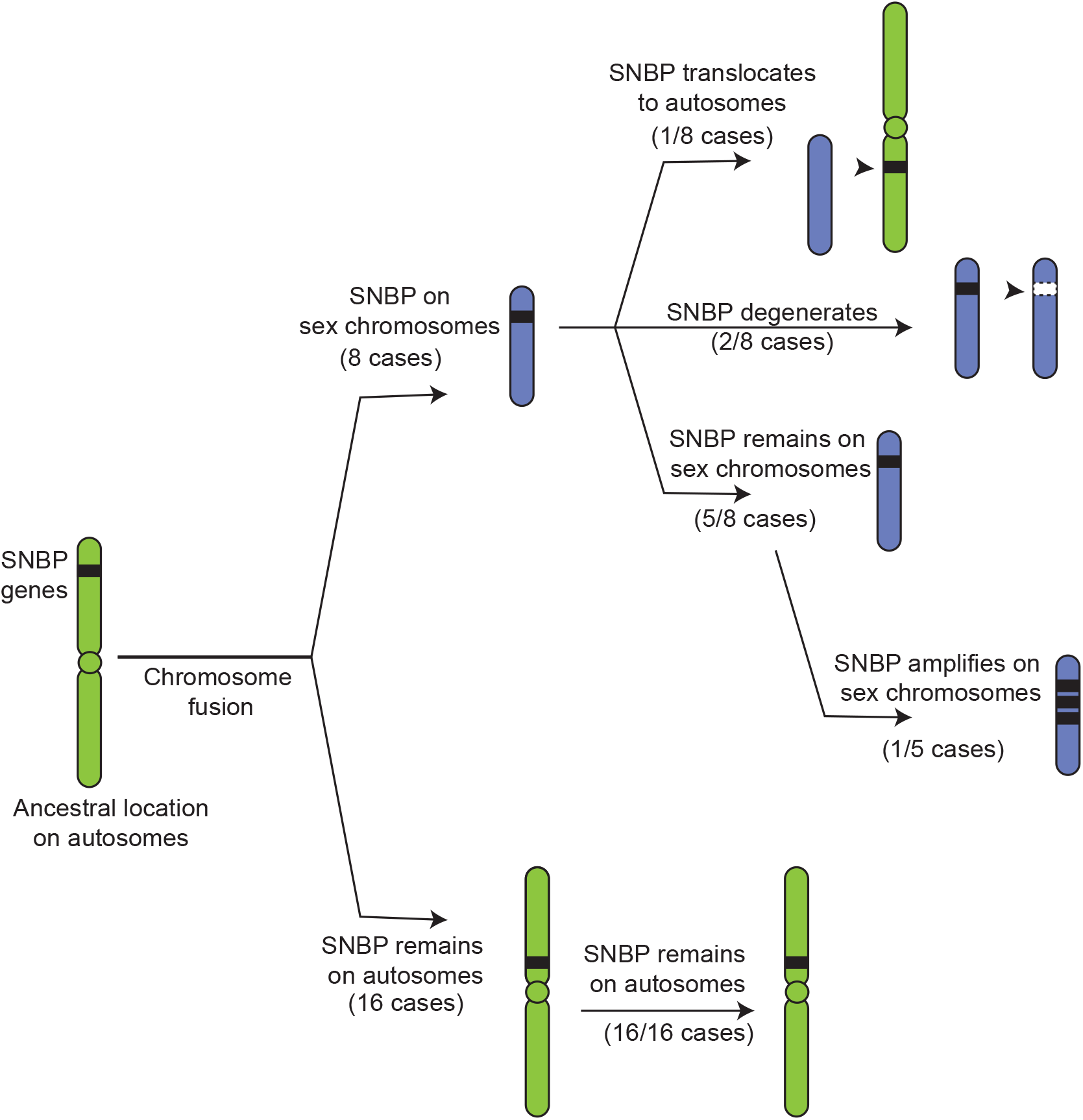
Evolutionary retention, degeneration or translocation of SNBP genes following chromosomal fusions. SNBP genes are ancestrally encoded on autosomes. Following chromosome fusion over *Drosophila* evolution, three SNBP genes, *CG14835*, *CG34269* and *ddbt* could become linked to sex chromosomes. We found that these SNBP genes translocated back to an autosome in 1/8 cases, degenerated in 2/8 cases, but were retained on neo-sex chromosomes in 5/8 cases. Among these five cases, we observed one amplification event– *ddbt* amplified to six copies in *D. repletoides.* In contrast, other SNBP genes that remained autosomal were strictly retained in 16/16 cases. These retention patterns differ significantly between sex chromosomes and autosomes (Fisher’s exact test, P=0.03). We also found in other species, except the *montium* group species, these three genes are widely conserved (one loss of *CG14835* and *CG34269* and two losses of *ddbt*).

### Loss of SNBP genes in the *montium* group coincides with X-Y chromosomal fusion

Our phylogenomic analyses also highlighted one *Drosophila* clade— the *montium* group of species (including *D. kikkawai*)— which suffered a precipitous loss of at least five SNBP genes that are otherwise conserved in sister and outgroup species (Fig 3). A closer examination allowed us to infer that six different SNBP genes underwent 11 independent degeneration events in the *montium* group (Fig 5A). Intriguingly, five of six SNBP genes lost in the *montium* clade (*Mst77F, Prtl99C, Mst33A, tHMG, CG14835*) are also among the eight SNBP genes subject to sex chromosome-specific amplifications in other *Drosophila* species (Fig 3 and Table 2). Notably, we did not find SNBP amplification events in any species of the *montium* clade. Given our hypothesis that autosomal SNBP genes might be linked to the suppression of meiotic drive (above), we speculated that the loss of these genes in the *montium* group of *Drosophila* species may have coincided with reduced genetic conflicts between sex chromosomes in this clade.

How could such reduction in sex chromosomal genetic conflicts arise? An important clue came from a previous study, which showed that many ancestrally Y-linked genes are present in females because of possible relocation to autosomes in the *montium* group [55]. We revisited this question to pinpoint which Y chromosomal gene translocations coincided with SNBP degeneration in this lineage. Using the available assemblies with Illumina-based chromosome assignment, we surprisingly found that most ancestrally Y-linked genes are not linked to autosomes as was previously suggested (Fig 6A) but instead are linked to the X-chromosome. Indeed, we were able to unambiguously infer X-chromosomal linkage for most ancestrally Y-linked genes in *D. kikkawai* (7/10), *D. jambulina* (9/11), *D. bocqueti* (7/10), *D. aff. chauvacae* (7/8), and *D. triauraria* (11/12) (Fig 6A), although some ancestrally Y-linked genes are not assembled in the assemblies. Moreover, in *D. triauraria*, we found that eleven of twelve ancestrally Y-linked genes, *i.e.,* all except *JYalpha,* are located on the same region of the X chromosome (Fig 6B). The most parsimonious explanation for these findings is a single translocation of most of the Y chromosome to the X chromosome via a chromosome fusion in the ancestor of the *montium* group of species. Afterward, some of these genes relocated back to the Y chromosome in some species (Fig S6; Supplementary text).

**Figure 6.**
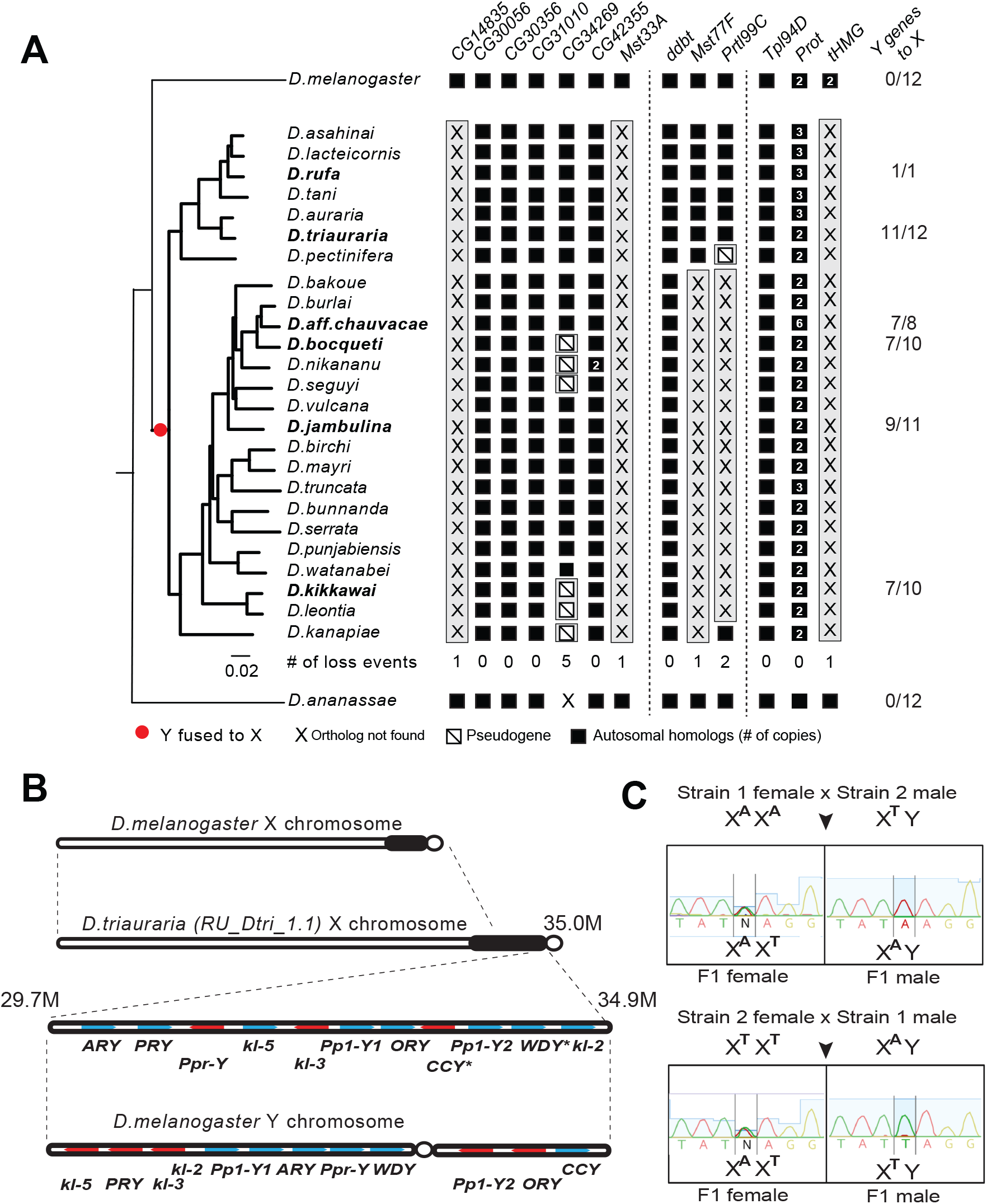
A dramatic loss of SNBP genes coincided with a fusion of X and Y chromosomes in the *montium* group species. **(A)** Using a phylogeny of species from the *montium* group, we traced the retention or loss of SNBP genes that are otherwise primarily conserved across other *Drosophila* species. Genes retained in autosomal syntenic locations are indicated in black squares, whereas pseudogenes are indicated by an empty square with a diagonal line. We traced a total of 11 independent pseudogenization events. Three of these pseudogenization events occurred early such that all species from this group are missing intact *CG14835, Mst33A* and *tHMG*. Three other SNBP genes were lost later (in some cases on multiple occasions) and are, therefore, missing only in a subset of species. For example, we infer that *CG34629* was lost on at least five independent occasions (and also in outgroup species *D. ananassae*). We correlated this dramatic loss of otherwise-conserved SNBP genes with the X-chromosome linkage of genes that are ancestrally Y-linked in other *Drosophila* species, shown on the right. Our ability to trace chromosomal linkage varied across different species. We traced 12 genes that are on Y chromosomes in most related species, including *D. melanogaster* and *D. ananassae*. Although some previously Y-linked genes were not assembled in current assemblies, we found that the most are now X-linked (*e.g.,* 11/12 in *D. triauraria*, 9/11 in *D. jambulina*, and 7/10 in *D. bocqueti* and *D. kikkawai*). **(B)** We traced the chromosomal arrangement and linkage of ancestrally Y-linked genes in *D. triauraria* using genome assemblies and genetic crosses in (C). We were able to show that the *D. triauraria* X chromosome represents a fusion of the X chromosome (*e.g.,* from *D. melanogaster*) and chromosomal segments containing 11 protein-coding genes that are typically found on the Y chromosome (*e.g.*, from *D. melanogaster*). Genetic crosses confirmed the X-linkage of 9 of these previously Y-linked genes, but the lack of allelic differences in *D. triauraria* prevented us from confirming this for two genes: *CCY* and *WDY*. **(C)** An example of the genetic cross used to verify X-linkage. Using genetic crosses between different *D. triauraria* strains that have allelic variation in Y-linked genes, we evaluated whether male flies inherit alleles of these genes maternally, paternally, or from both parents. We observed only maternal inheritance, confirming the X-chromosomal linkage of these genes.

We carried out genetic mapping studies to confirm our unexpected inference of a Y-to-X translocation for the *montium* group. For this, we performed genetic crosses between different *D. triauraria* strains harboring polymorphisms on ancestrally Y-linked genes [56]. To unambiguously infer the chromosomal location of these genes, we focused on whether SNPs in these genes were paternally or maternally inherited. If the SNPs were Y-linked, we would expect them to be strictly paternally inherited whereas we would expect both paternal and maternal inheritance in the case of autosomal linkage. In contrast to these expectations, all F1 males inherit alleles of ancestral Y-linked genes from their mother, unambiguously indicating their X-chromosomal linkage (Fig 6C).

The translocation of a large segment of the Y-chromosome to the X-chromosome in the *montium* group would render any X-versus-Y meiotic drive encoded in this chromosomal region obsolete or costly. As a result, there would be active selection to jettison such meiotic drive systems on both the X and Y chromosomes. Indeed, no meiotic drive has been documented in the *montium* species even though it is rampant in many other *Drosophila* lineages [38]. Subsequently, any autosomal SNBP genes required to suppress meiotic drive would become dispensable, leading to their degeneration. Thus, our hypothesis of genetic conflicts between sex chromosomes can also explain the loss or degeneration of SNBP genes following alleviation of the sex-chromosomal conflict as well (Fig 7).

**Figure 7.**
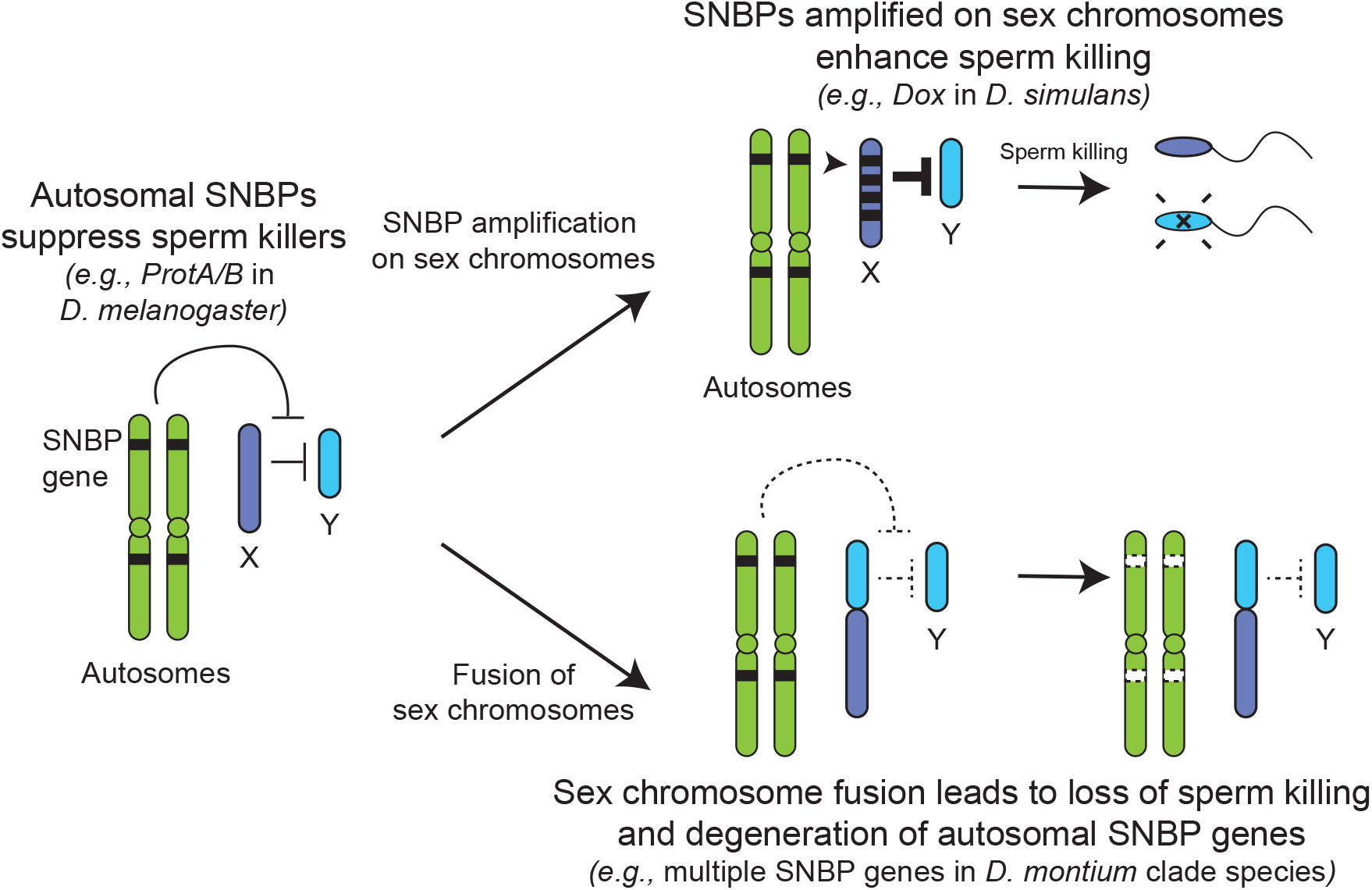
Genetic conflict between sex chromosomes may explain the rapid turnover of SNBP genes in *Drosophila* species. SNBP genes are ancestrally encoded on autosomes where we hypothesize that some of them act to suppress meiotic drive between sex chromosomes (*e.g., ProtA/B*). However, in some cases, paralogs of these SNBP genes duplicate onto sex chromosomes where they undergo dramatic amplification. We propose that this amplification creates an opportunity for them to act as meiotic drive elements themselves (*e.g., Dox*), imbuing sex chromosomes that inherit them with transmission advantages. A fusion of the sex chromosomes (*e.g., D. montium* species group) leads to a loss of meiotic competition between sex chromosomes, which will subsequently lead to the loss or degeneration of the suppressing SNBP genes on autosomes since their drive suppression functions are rendered superfluous.

## Discussion

Our analyses of *Drosophila* SNBP genes reveal many similar patterns to rapid protamine evolution observed in mammals, including a pervasive pattern of positive selection. However, there are also some dramatic differences, which may stem from distinct biological functions and selective pressures. One of the most dramatic differences is the evolutionary turnover of SNBP genes in *Drosophila* versus mammals. Most mammals share four SNBP genes: *TNP1, TNP2, PRM1,* and *PRM2*. In contrast, SNBP repertoires vary extensively between *Drosophila* species, with both dramatic gains and losses. We discovered several independent, species-specific amplifications of SNBP genes from 5 to >50 copies that preferentially occurred on sex chromosomes (Fig 3 and Table 2). Conversely, we also found that several SNBP genes were lost in the *montium* group, coinciding with an X-Y chromosome fusion event (Fig 6 and Table 2). Moreover, in other lineages, three SNBP genes that became linked to sex chromosomes via chromosomal fusions either degenerated or translocated back to autosomes (Fig 5). We hypothesize that many SNBP genes, such as the *ProtA/B* genes in *D. melanogaster*, arise and are retained on autosomes to act as suppressors of meiotic drive [39]. In contrast, sex chromosomal amplifications of SNBP genes, like the *Dox* amplification, act as meiotic drivers, potentially by disrupting the transcription or translation of autosomal SNBP genes or disrupting the histone-to-protamine transition via their protein products [36, 37, 40]. X-Y chromosome fusions eliminate the extent of meiotic drive and may lead to the degeneration of otherwise conserved SNBP genes, whose functions as drive suppressors are no longer required. Thus, unlike in mammals, sex chromosome-associated meiotic drive appears to be the primary cause of SNBP evolutionary turnover in *Drosophila* species.

Why would meiotic drive only influence *Drosophila*, but not mammalian, SNBP evolution? One important distinction may arise from the timing of SNBP transcription. In *D. melanogaster,* SNBP genes are transcribed before meiosis but translated after meiosis [29, 43, 57]. Thus, SNBP transcripts from a single allele, *e.g.*, X-linked allele, are inherited and translated by all sperm, regardless of which chromosomes they carry. Consequently, they can act as meiotic drivers by causing chromatin dysfunction in sperm without the allele, *e.g.*, Y-bearing sperm. In contrast, mammalian SNBPs are only expressed post-meiosis [58, 59] and therefore individual SNBP alleles can only affect the chromatin states of sperm in which they reside. Therefore, mammalian SNBP genes are not likely to evolve as meiotic drivers.

Our findings do not rule out the possibility that forces other than meiotic drive are also important for driving the rapid evolution and turnover of SNBP genes in *Drosophila* species. Indeed, we note that only some SNBP genes undergo both amplifications and losses across *Drosophila* species (Fig 3 and Table 2). Other SNBP genes that do not experience such dramatic turnover still evolve rapidly (Table 2). Moreover, many SNBP genes in *D. serrata*, a *montium* group species with a fused X-Y chromosome, continue to evolve under positive selection (Table 1). Their rapid evolution may be driven by sperm competition, just as in mammals.

Another possible selective pressure on SNBP genes may come from endosymbiotic *Wolbachia* bacteria. One means by which *Wolbachia* manipulate *Drosophila* hosts to ensure their preferential propagation is by mediating cytoplasmic incompatibility, in which embryos produced between infected males and uninfected females fail to de-compact the paternal pronucleus and arrest in development [60, 61]. Recent studies have revealed that *Wolbachia* produces toxins that prevent the deposition of SNBP during spermatogenesis and delay SNBP removal in embryos post-fertilization to accomplish this cytoplasmic incompatibility [62, 63]. Indeed, deletions of *ProtA/ProtB* exacerbate the intensity of cytoplasmic incompatibility imposed by *Wolbachia* in *D. melanogaster* [64]. Thus, selective pressure from *Wolbachia* toxins could also provide the selective pressures for SNBP innovation.

Our analyses have focused on SNBP genes that can be readily identified because they possess HMG DNA-binding domains. However, this does not represent a comprehensive list of SNBP genes. Indeed, recent findings have shown that the *atlas* SNBP gene from *D. melanogaster* does not encode an HMG domain entirely and has arisen *de novo* [33]. Such genes might use an alternate means to bind DNA or may indirectly affect the localization of other SNBP proteins and therefore sperm chromatin. However, even focusing on the HMG-domain containing SNBP genes reveals an unexpected relationship between their essential function and evolution. SNBP genes encoding essential functions arose recently and have been frequently lost in *Drosophila* species. In contrast, SNBP genes less essential for male fertility are prone to evolve under positive selection. This suggests that SNBP genes with redundant roles in male fertility are more likely to acquire new function.

One clue for this unexpected lability emerges from discovering SNBP genes that are essential for *D. melanogaster* fertility but were lost in other species. For example, *ddbt* is a conserved SNBP gene with essential sperm telomere-capping function in *D. melanogaster*; loss of *ddbt* leads to defects in telomere capping and induction of telomeric fusions in embryos [31]. We find that the two lineages that lost *ddbt*— all *D. willistoni* subgroup species, and *D. albomicans* in the *D. immigrans* species group— each has a pair of neo-sex chromosomes due to the fusion of Muller element D and sex chromosomes [65–68]. Since the knockout of *ddbt* induces telomeric fusion events in *D. melanogaster* embryos, we hypothesize that the loss of *ddbt* might have led to these chromosome fusions that cause the independently evolved neo-sex chromosomes. Alternatively, since telomeric sequences rapidly evolve in an ‘arms-race’-like dynamic with telomere-binding proteins across *Drosophila* species [69], particular chromatin rearrangements may have obviated the essential function of *ddbt* in some species, as has been hypothesized for other paternal-effect lethal chromatin genes [70]. This suggests that the nature of essential SNBP functions can be idiosyncratic and species-specific, arising from the underlying rapid evolution of chromatin, as has been hypothesized for evolutionarily young but essential genes involved in centromere and heterochromatin function [50, 71].

## Materials & Methods

### Molecular evolutionary analyses of SNBP genes

We use PAML 4.9 [48] to calculate protein evolution rates (dN/dS) of SNBP genes and conduct site-model, branch-model and branch-site model analyses. We compare the protein evolution rates of SNBP genes to the genome-wide rates (8521 genes) from the 12 *Drosophila* genomes project [72]. Many SNBP genes were missed in previous analyses because they are rapidly evolving and hard to align. Therefore, we aligned the coding sequences of orthologous SNBP genes from the same six *Drosophila* used to calculate dN/dS previously [72], and only used the orthologous sequences. For all phylogenic analyses, we first constructed maximum-likelihood trees using iqtree 2.1.3 [73] using parameters “-m MFP -nt AUTO -alrt 1000 -bb 1000 -bnni”. We then calculated protein evolution rates of SNBP genes using the same parameters with the generated gene trees (model = 0 and CodonFreq = 2).

For the site-model codeml analyses, we analyzed 9–17 unambiguous orthologs from species in the *melanogaster* group to increase the power and accuracy. We compared NS sites models M1a to M2a, and M7 or M8a, to M8 using likelihood ratio tests to ask whether models allowing a class of sites where dN/dS exceeds 1 (*e.g.,* M8) provide a better fit to the data than models that disallow positive selection (*e.g.,* M7, M8a). We also used different codon parameter values (CodonFreq = 0, 2, 3) in our analyses to check whether our results were robust.

For the branch-model, we first simplified the tree by reconstructing the ancestral sequences of ampliconic genes in each species using MEGAX (10.1.8). We compared models by assigning different protein evolution rates to branches using PAML (CodonFreq = 2) and likelihood ratio tests. Our models include the null model (same protein evolution rate across branches; model = 0), all X-linked branches share different protein evolution other than other branches, and the early duplication branches on both the X-linked copies and the parental copy share different protein evolution other than other branches (model = 2). The model with a higher protein evolution rate at early duplication branches fits best across all models, so we applied this setting to conduct the branch-site test. We compared two models with all sites share the same protein evolution rate (fix_omega = 0) and various evolution rates (fix_omega = 1).

To look for positive selection in two individual lineages, *D. melanogaster* and *D. serrata*, we applied McDonald–Kreitman tests to compare within-species polymorphism and between-species divergence [74]. We used *D. simulans* as the closely related species for the *D. melanogaster* analysis, and *D. bunnanda* for the *D. serrata* analysis. To further polarize the changes in each lineage, we inferred ancestral sequences of *D. melanogaster* and *D. simulans* using seven species in the *D. melanogaster* group (*D. melanogaster*, *D. simulans, D. mauritiana, D. sechellia, D. yakuba, D. erecta,* and one other well-aligned outgroup species (*D. biarmipes, D. elegans, D. eugracilis, D. ficusphila, or D. rhopaloa*) using MEGAX (10.1.8) [75]. Similarly, we used five species in the *montium* group (*D. serrata, D. bunnanda, D. birchii*, *D. truncata* and *D. mayri*) to polarize the *D. serrata*-specific changes. We extracted population data from public datasets of >1000 *D. melanogaster* strains [76, 77] and 111 *D. serrata* strains [46]. We conducted both polarized and unpolarized McDonald–Kreitman tests using R scripts (https://github.com/jayoung/MKtests_JY) and confirmed our findings using an online server ([78]; http://mkt.uab.es/mkt/MKT.asp).

### Searching for homologs of SNBP genes in *Drosophila* and outgroup species

We used tblastn and reciprocal blastx to search homologs of all SNBP genes across all genome assemblies [53] using *D. melanogaster* protein sequences as queries. Since SNBP genes are rapidly evolving, we used the following parameters: e-value < 1e-2, amino acid identity > 20% and blast score > 10. We further required that the best reciprocal blastx hit when searching *D. melanogaster* genes was the original query gene, to ensure that we were recovering true orthologs. To further confirm questionable orthologs, *e.g.,* only one species with the homolog in the lineage, we examined its synteny (conserved genomic neighboring genes), anticipating that orthologs should also have shared syntenic contexts. We also examined the syntenic regions of SNBP genes and conducted blastx using a lower threshold (e-value < 1) to confirm the loss of SNBP genes in some lineages, especially in the *montium* group species.

We used the abSENSE package (https://github.com/caraweisman/abSENSE; [45]) to calculate the probability of not detecting homologs in more diverged species (using E-value=1), enabling us to distinguish whether our inability to detect homologs was due to rapid divergence or true absence.

### Transcriptomic analyses of SNBP genes

We combined the public gene annotations with our own annotated SNBP gene annotations and mapped publicly available transcriptome datasets (Table S12) to the genome assemblies using HiSAT2 (v2.2.1 with parameters –exon and –ss to specify the exon positions and splice sites; [79]). We then estimated the expression levels using the gene annotations as input for stringtie (v2.1.4 with parameters -dta -G to specify annotation files; [80]).

### Assigning sex-linkage of contigs in genome assemblies

We mapped Nanopore and Illumina reads from male samples (sometimes from mixed-sex or female samples) to genome assemblies using minimap2 [81] and bwa-mem [82] using the default parameters. We calculated coverage of each site using samtools depth and estimated the median coverage of each 10-kb window and each contig. We then examined the genome-wide distribution of 10-kb window coverage using the density function in R. We called two peaks of coverage using turnpoints function in R. As we expected, the prominent peak with higher coverage mostly represents autosomal regions whereas the lower coverage peak mostly represents sex-linked regions. We used the average of coverage from the two peaks as the threshold to assign autosomal and sex-chromosomal contigs.

We also confirmed our assignments using the sequence-homology method. We identified orthologs of 3285 highly conserved genes in all species using BUSCO v5.0.0 with default setting and diptera_odb10 database [54]. Since the X chromosome is conserved across *Drosophila* species, we identify orthologs of X-linked genes to assign X-linked contigs. Lastly, for species with Illumina data from both males and females, we distinguished X-from Y-chromosome sex-linked contigs using a previously described method that assigns using a previously described method [83]. We could not reliably distinguish their sex chromosomes from autosomes in *Lordiphosa* species (probably due to high heterozygosity or lower assembly quality) and therefore could not confidently assign the chromosomal location of their SNBP amplifications.

We used genetic mapping to examine the X-linkage of ancestral Y-linked genes in *D. triauraria*. We first identified polymorphic SNPs in *D. triauraria* SNBP genes using publicly available Illumina data [56] and called SNPs (bcftools call -m -Oz) in each strain [84, 85], and took advantage of the fact that F1 males will only inherit maternal X alleles. We crossed two strains with different alleles of ancestrally Y-linked genes and genotyped these genes in F1 males using PCR (Table S13) and Sanger sequencing to examine whether their allele-specific inheritance patterns were more consistent with Y-linkage (paternal inheritance), X-linkage (maternal inheritance) or autosomal linkage (both paternal and maternal inheritance).

### CRISPR/Cas9 knockout and fertility assays

To generate the *CG30056* knockout strain, we first cloned two guide RNAs, targeting either the 5’ or 3’ end of CG30056, into pCFD4-U6:1_U6:3tandemgRNAs (Addgene #49411) using NEBuilder® HiFi DNA Assembly Master Mix (NEB catalog E2621). For the repairing construct, we used independent PCR reactions to amplify 3xP3 DsRed, the backbone of pDsRed-attP (Addgene #51019) and ∼1kb homologous sequences of upstream and downstream of *CG30056* using PCR independently. These four fragments were annealed using NEBuilder® HiFi DNA Assembly Master Mix. We then used the Q5® Site-Directed Mutagenesis Kit (NEB catalog E0554S) to mutate the gRNA target PAM sites on the repairing construct so that they would not also be targeted by the guide RNAs we used. All primers are listed in (Table S13). These two constructs were injected into y[1] M{GFP[E.3xP3]=vas-Cas9.RFP-}ZH-2A w[1118] (BDSC #55821) embryos by BestGene. The resulting transgenic flies were backcrossed to *yw*, selected using the DsRed marker, and balanced by *CyO*. The transgenic flies were further confirmed by PCR using independent primer sets (Fig 3A and Table S13).

For comparisons, we also tested male fertility effects of two SNBP genes previously shown to be essential (*Mst77F*) or not (*Tpl94D*) for male fertility. We obtained *Mst77F* (Δ1) knockout flies (kindly provided by Dr. Benjamin Loppin [28]) and two fly strains carrying large deletions spanning *Mst77F* (BDSC #24956 and 27369). For *Tpl94D*, we used a mutant from the *Drosophila* Gene Disruption Project (BDSC #26333; [86]) and a fly carrying a large deletion spanning this gene (BDSC #7672; [87]).

We obtained test males for fertility assays by crossing 2-5 day-old female virgin transgenic flies with other transgenic flies or flies carrying a large deletion (BDSC #30585). To measure their fertility, each resulting 2-5 day-old F1 male was crossed to two 2-5 day-old virgin females from the wildtype Oregon R *D. melanogaster* strain at room temperature. We transferred the mating pairs to new vials every three days and counted all resulting offspring from the first 12 days.

## Acknowledgements

We thank Drs. Cécile Courret, Janet Young, Phoebe Hsieh, Pravrutha Raman, and Isabel Mejia Natividad for comments on the manuscript and members of the Malik, Ahmad, and Henikoff labs and Drs. Barbara Wakimoto, Amanda Larracuente and John Sproul, for fruitful discussions. We thank Isabel Mejia Natividad for her help on the primer design, PCR and sequencing. We also thank Benjamin Loppin and the Bloomington *Drosophila* Stock Center for the *Drosophila* strains used. We are supported by a Damon-Runyon Cancer Research Foundation postdoctoral fellowship DRG 2438-21 (to C.-H.C) and National Institutes of Health grant R01-GM74108 (to H.S.M.). H.S.M is an Investigator of the Howard Hughes Medical Institute. This article is subject to HHMI’s Open Access to Publications policy. HHMI lab heads have previously granted a nonexclusive CC BY 4.0 license to the public and a sublicensable license to HHMI in their research articles. Pursuant to those licenses, the author-accepted manuscript of this article can be made freely available under a CC BY 4.0 license immediately upon publication.

## Supplementary Materials

**Figure S1.**
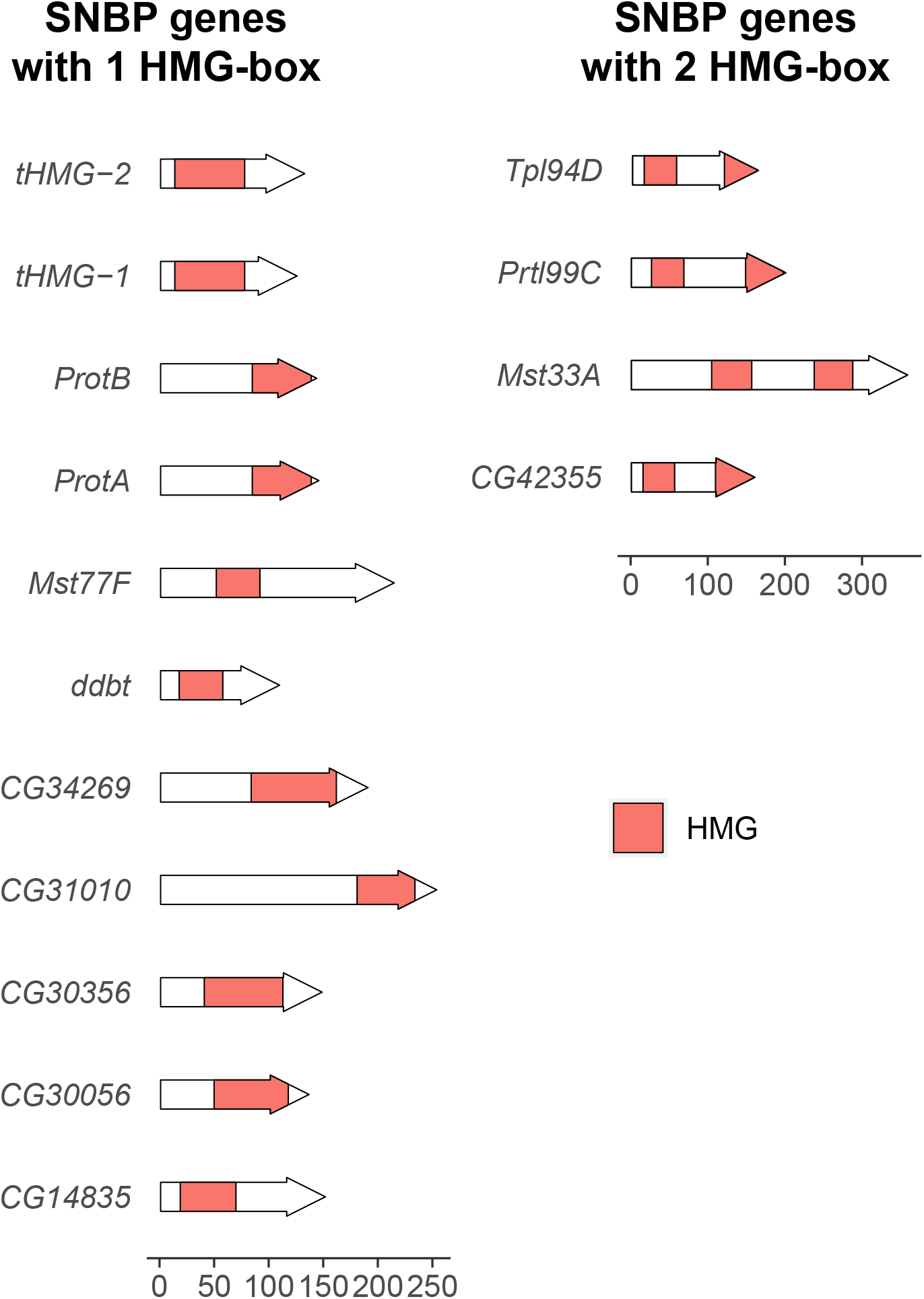
The number and position of high mobility group (HMG) boxes in *D. melanogaster* SNBP proteins. We plotted the location of HMG boxes in 15 SNBP proteins. Among them, 11 have only one HMG box and four of them have two HMG boxes. The position of HMG boxes varies between SNBP proteins. The scale is at the bottom of the figure.

**Figure S2.**
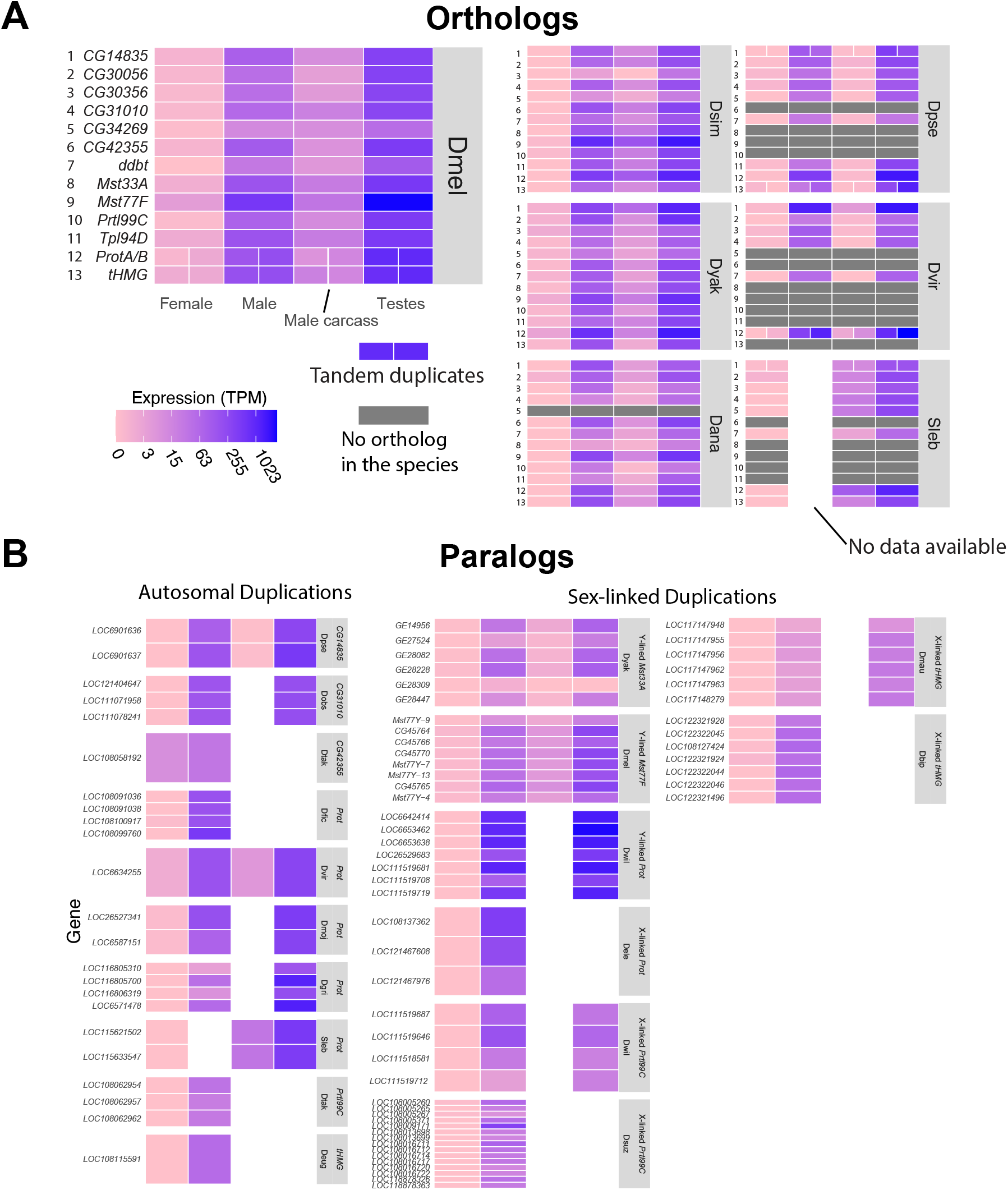
The expression level of SNBP genes (A) and their paralogs (B) from different tissues in *Drosophila* and *Scaptodrosophila* species. We estimated the expression of SNBP genes using publicly available transcriptome datasets (Table S12). We use colors to represent expression levels in each sample, and find almost all SNBP genes are expressed only in testes. The raw values are shown in Table S2.

**Figure S3.**
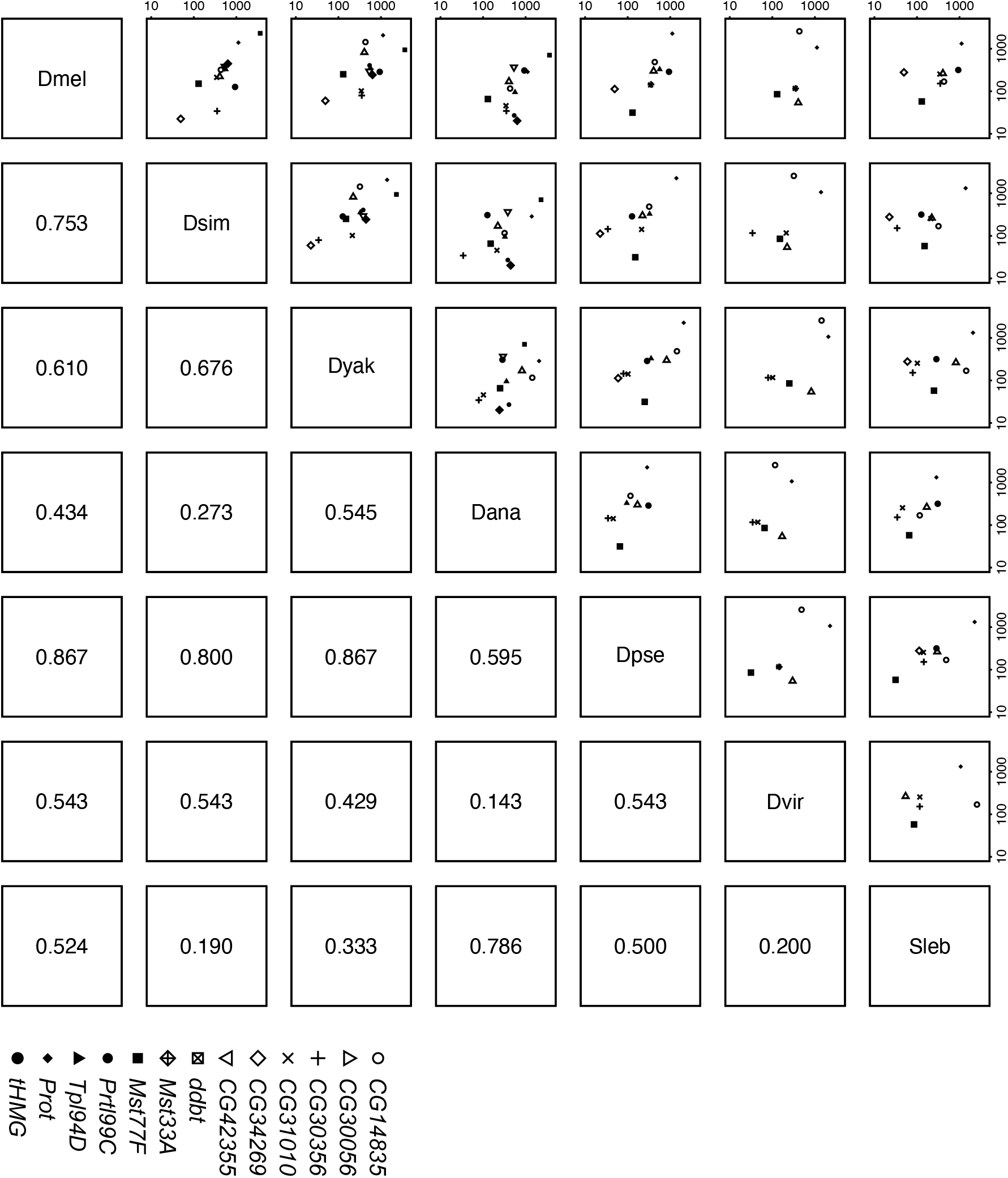
SNBP testis expression level is correlated across seven *Drosophila* species. We estimated the expression level of each SNBP gene in testes across species, and compared the relative expression level of orthologs to each other. The numbers below the diagonal are spearman rho. Our data suggest medium to high correlation between *Sophophora* species. The raw values are shown in Table S2.

**Figure S4.**
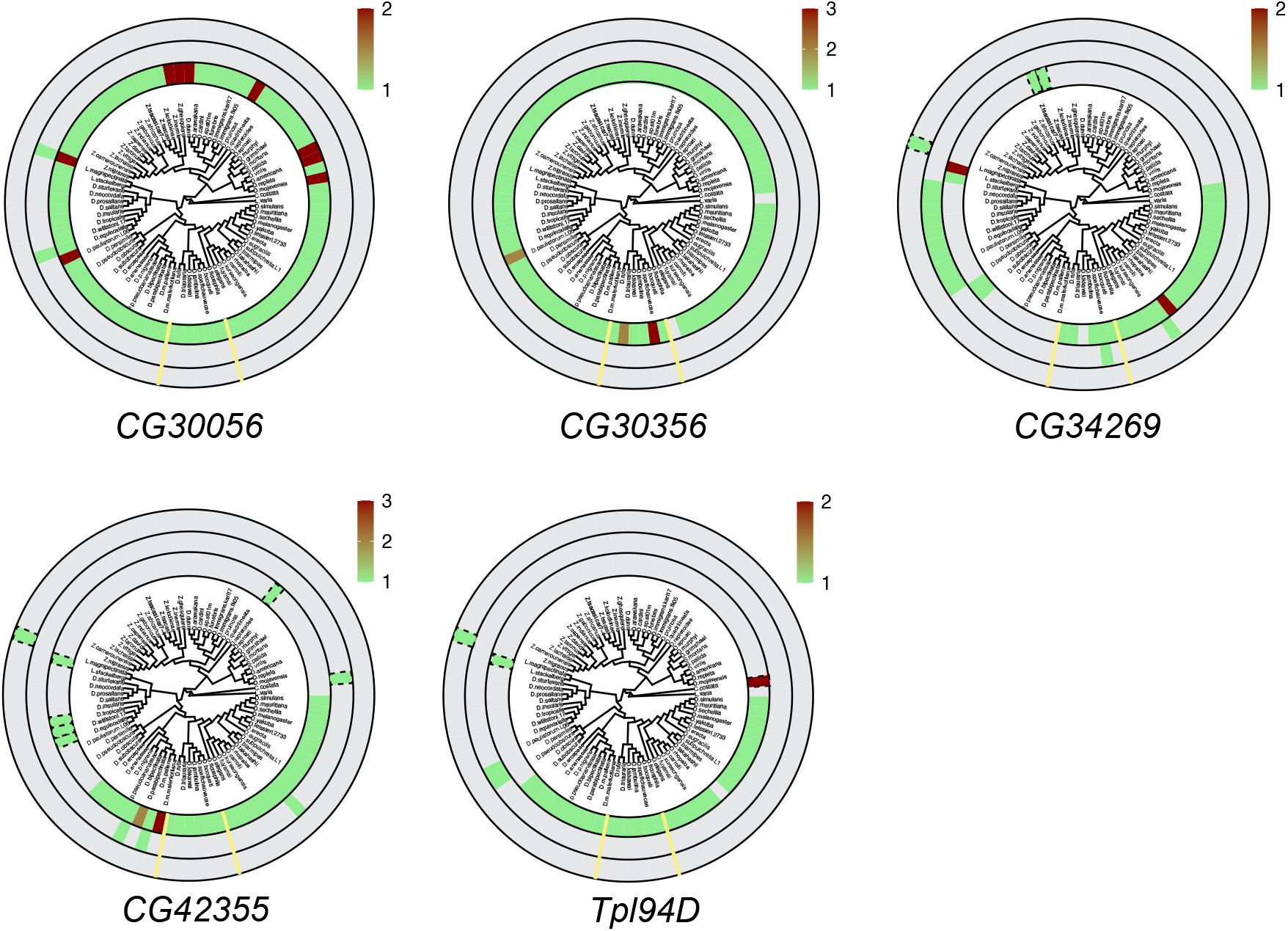
Copy numbers of six SNBP genes without gene amplification. We searched for homologs of each *D. melanogaster* SNBP gene in 78 distinct *Drosophila* species using reciprocal BLAST. We represent our findings using the same circular representation as Fig 3. The innermost ring indicates autosomal genes, the middle ring indicates sex-linked genes, and the outer ring shows genes with an ambiguous location.

**Figure S5.**
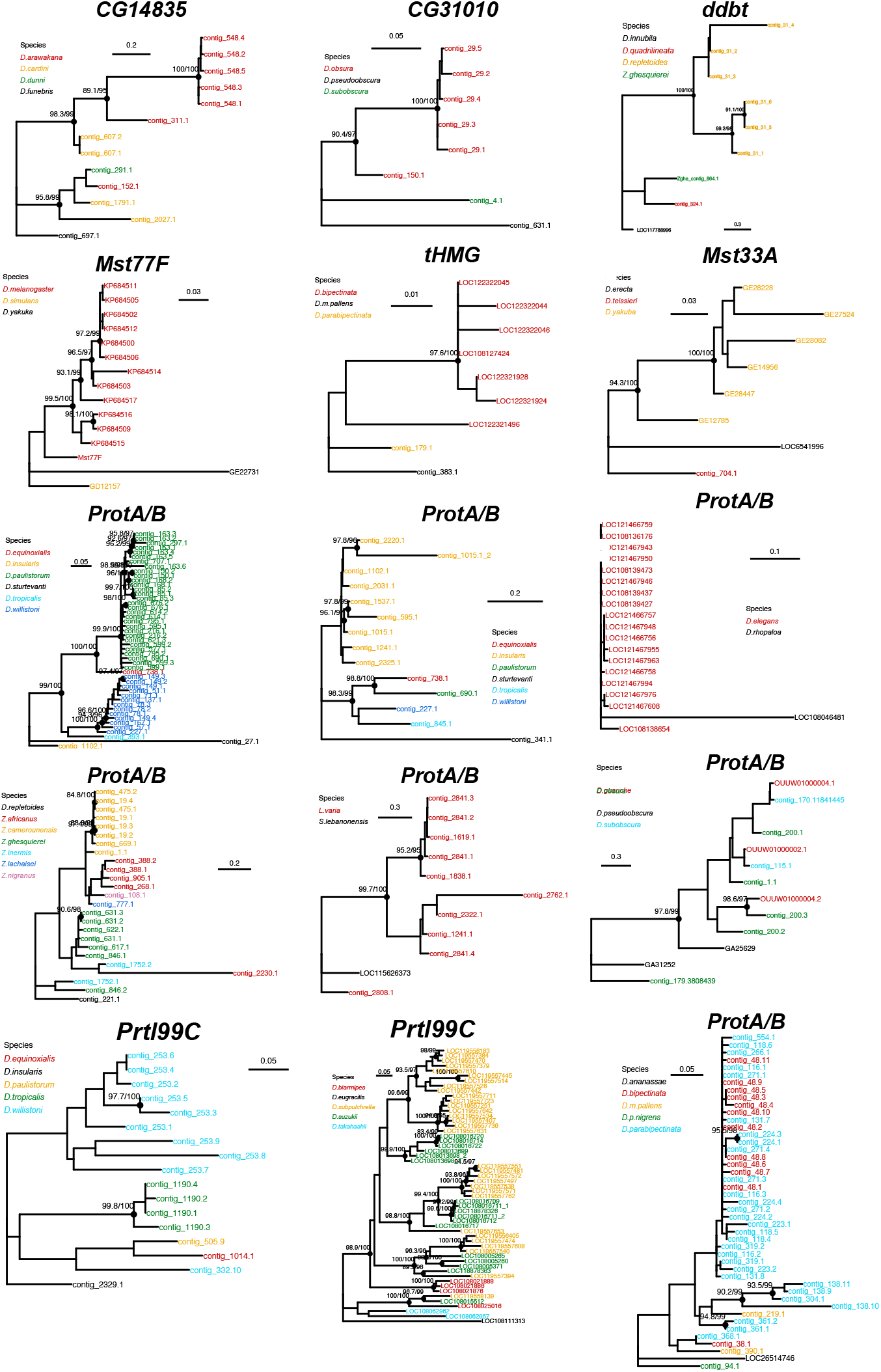
Phylogenies of amplified SNBP genes show recent duplications and concerted evolution. Phylogenetic analyses of the various SNBP genes with amplification demonstrate that most of these amplifications are evolutionarily young. The phylogeny also suggests concerted evolution among the amplified copies of *CG14835* in the *D. arawakana* clade and *Prtl99C* in the *D. suzukii* clade. The phylogenies from amplified copies of *tHMG* and *ProtA/B* in *Lordiphosa* species are not shown here because of the lack of good outgroup species and low-quality sequences.

**Figure S6.**
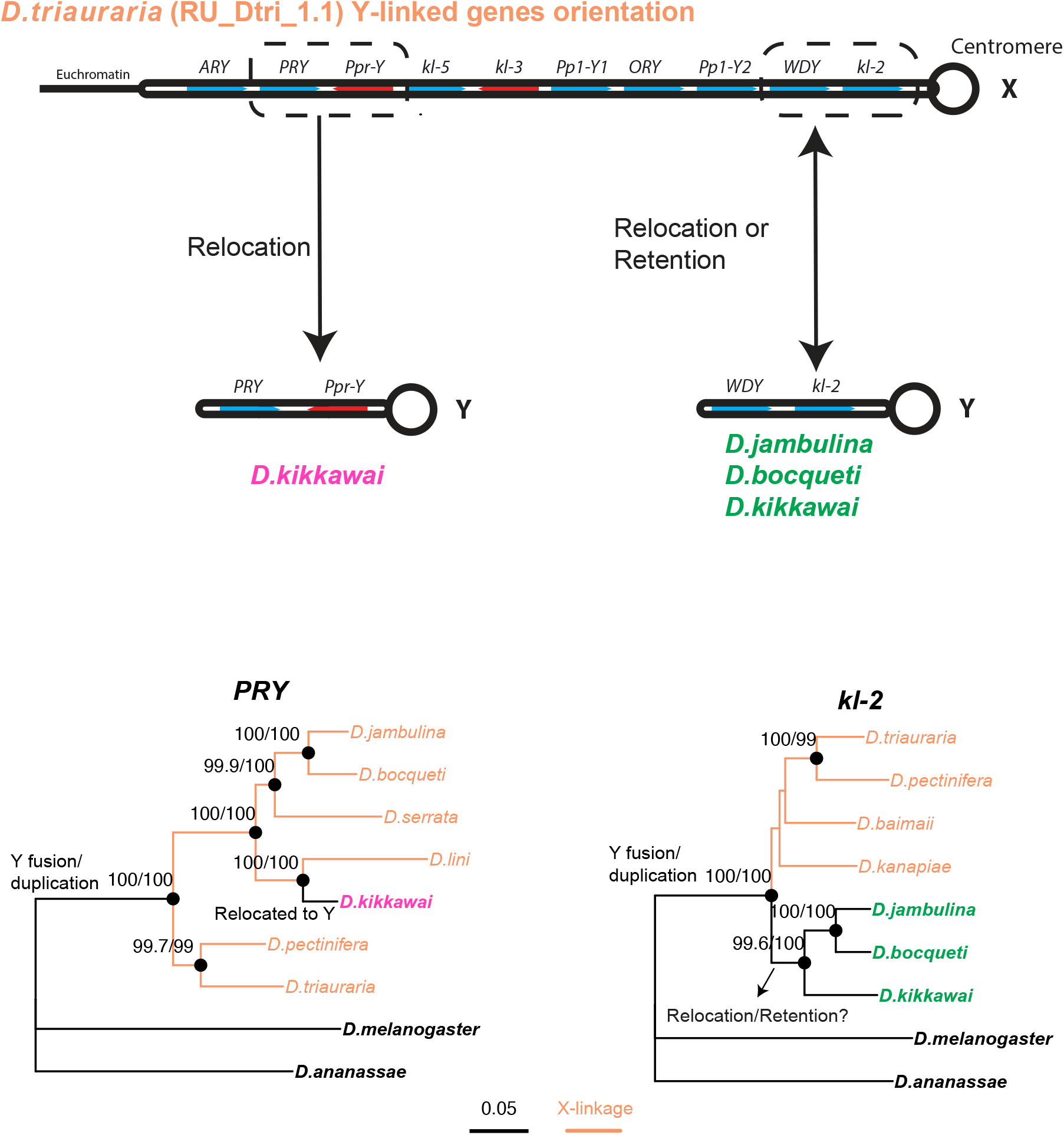
The gene phylogeny supports the secondary translocation of ancestrally Y-linked genes back to the Y chromosome. *PRY* is located on the same contig as *Ppr-Y*, and *WDY* is located on the same contig as *kl-2* in *D. kikkawai*. The neighboring arrangement of these two genes is conserved between *D. kikkawai* and *D. triauraria* even though they are located on different chromosomes in the two species. Both genes are consistent with the species tree in the species we have sequence information. Phylogenetic analyses of *PRY* suggest that it relocated back to the Y chromosome from the X chromosome in *D. kikkawai* lineage. In contrast, we cannot infer whether the Y-linked *kl-2* and *WDY* of *D. kikkawai* are derived from an X-linked ancestor or are located on Y ancestrally.

**Table S1. The probabilities of homologs of young SNBP genes being undetected in each species, estimated using abSENSE.** abSENSE used the blast scores in related species with detected orthologs and calculated the Blast scores of the orthologs in more related species given the divergence of species. Then it estimated the probability of failing to detect a homolog (if it were present) in species of various divergence levels (using E-value=1).

**Table S2. The expression level (TPM) of SNBP genes in different tissues of Drosophila and Scaptodrosophila species.** We estimated the expression levels of SNBP genes using publicly available transcriptome datasets (Table S12). The data is also illustrated in Fig S3.

**Table S3. The basic sequence information of SNBPs and their homologs in each species.** We collected the sequences of SNBPs and their homologs from the NCBI database. We calculated the isoelectric point and length of each protein using Geneious 2022.1.1 (https://www.geneious.com).

**Table S4. Evolutionary rates of SNBP genes and other genes in D. melanogaster subgroup species.** We used PAML to estimate evolutionary rates of SNBP genes using the same parameters and the same six *Drosophila s*pecies used in the 12 *Drosophila* genomes project [72]. The evolutionary rates of other genes were acquired from the 12 *Drosophila* genomes project [72].

**Table S5. McDonald–Kreitman test results for SNBP genes in *D. melanogaster* and *D. serrata*.** We looked for positive selection in two lineages, *D. melanogaster* and *D. serrata,* using McDonald–Kreitman tests to compare within-species polymorphism to between species divergence [74]. We used *D. simulans* as the closely-related species for the *D. melanogaster* analysis, and *D. bunnanda* for the *D. serrata* analysis.

**Table S6. No evidence for positive selection on SNBP genes using site model in PAML in *D. melanogaster* subgroup species.** We aligned 9–17 unambiguous orthologs from species in the *D. melanogaster* group to test whether a subset of sites evolves under positive selection. We compared NSsites models M1a to M2a, and M7 or M8a to M8 using likelihood ratio tests. We ran each model using several codon parameter choices (CodonFreq =0, 2, 3) to check whether results were robust. For example, *CG30056* shows signal of difference selection strength across site using CodonFreq =2 (p=0.0003), but not CodonFreq=0 or 3 (p=1 and 0.16 respectively).

**Table S7. Low frequency of polymorphic mutations with severe effects on SNBP genes in D. melanogaster population.** We extracted population data using an available dataset of >1000 *D. melanogaster* strains [76, 77], and documented changes that might inactivate SNBP proteins.

**Table S8. Chromosomal assignments for each contig containing SNBP genes.** We assigned the location of SNBP-containing contigs using synteny (Muller elements) and coverage analysis. We used BUSCO genes on these contigs to assign their most likely Muller element. We also mapped available male Nanopore or Illumina reads to the assemblies and estimated coverage on the contigs compared to autosomal contigs. If the normalized read coverage is significantly less than 1, we assign the contigs to either X or Y chromosome.

**Table S9. The copy number and chromosome location of SNBP homologs using BLAST in each species.** We summarized the data from Table S8 and also manually curated data from some amplified SNBP genes using extra assemblies or Illumina reads (shown in red). To determine the chromosomal location of some amplified SNBP genes, we mapped male and female Illumina reads from different resources to the assemblies of 10 species (Table S11). This allows us to assign contigs to X or Y chromosome unambiguously. For *D. melanogaster* and *D. simulans*, we used assemblies with better contiguities (GCA_000778455.1[89] and GCA_004382185.1 [90]).

**Table S10. The PAML analyses reveal different selection forces in *tHMG* duplicates of *D. simulans* clade species.** We analyzed *tHMG* copies from *D. simulans* clade species to infer their selective pressures (Fig. 4). We compared branches with different protein evolution rates using likelihood ratio tests (CodonFreq =2). Our models include the null model (same protein evolution rate across branches), a model where all X-chromosome branches share a rate that is different from the rate on all other branches (“all X”), and a model where the early duplication branches on both the X-linked copies and the parental copy share a rate that is different from all other branches (“Duplication”). We compared two models with all sites that share the same protein evolution rate (fix_omega = 0) and various evolution rates (fix_omega = 1), and did not find the evidence of positive selection. The duplication model fits best across all models, so we also used this model to conduct a branch-site test. No evidence for positive selection was using the branch-site test.

**Table S11. The location and degeneration of SNBP genes in species with neo-sex chromosomes.** Chromosomal locations of each SNBP gene in species with neo-sex chromosomes. The data is illustrated in Figure 5.

**Table S12. Sequence data resources and information used in this study**

**Table S13. Primer sequences used in this study.**

## Supplementary text

### Low frequency of impaired mutation in all SNBP genes

We next investigated whether the essentiality of SNBP genes correlated with their evolutionary age or retention. We first investigated the intra-species retention of all 15 SNBP genes, taking advantage of extensive sequencing of *D. melanogaster* strains by many labs [1, 2-10]. We found that loss-of-function variants of SNBP genes segregate at very low frequencies (< 1%) among *D. melanogaster* strains (Table S7). The only exceptions are *CG14835* (1.5% frequency of frameshift mutation in worldwide populations) and *ddbt* (1.2% frequency of loss of start codon in non-African populations). However, the *ddbt* mutation is likely to be benign owing to an alternate start codon just a few codons downstream of the canonical start site. In contrast, variants that are not likely to impair function (small in-frame indels) can segregate at higher frequency, *e.g.,* a 15-bp insertion variant of *tHMG2* is present at 70% frequency in worldwide *D. melanogaster* populations. This suggests nearly strict retention of all SNBP genes, whether they were shown to be essential for male fertility in laboratory experiments or not, in all sequenced strains of *D. melanogaster*.

### The amplification of *tHMG* copies in *D. simulans* clade species

Based on evolutionary reconstruction, we infer that a gene duplication of the ancestral, autosomal *tHMG* first arose in the euchromatin of the X chromosome, proximal to *CG12691*. A subsequent duplication spanning X-linked *tHMG* and part of *CG12691* arose and then tandemly duplicated in the heterochromatic region of the X chromosome, close to the *flamenco* locus. As a result of this tandem duplication, the amplicon consisting of a *tHMG* and *CG12691* fragment is present in more than 15 copies on a heterochromatic region of the X chromosome in *D. simulans* and *D. mauritiana,* but was lost in *D. sechellia*, at least in the sequenced strains. In addition to this X chromosomal expansion, we also found a few degenerated copies of *tHMG* on the 3R heterochromatic region and the Y chromosome (Fig 4A). We also found that copies from the X-linked heterochromatic region are highly homogeneous within species, but diverged between species (Fig 4C). Such a pattern of concerted evolution, in which paralogous SNBP genes from species are more similar to each other than to homologs from other species [11], is also seen in other cases of SNBP amplifications (Fig S5). We note that we detected different copy numbers (all more than 30) of *tHMG-hetX* across three sequenced strains of *D. simulans* we surveyed. This difference is due to both the incomplete assemblies of this region and strain differences. Surprisingly, we found that the new X-linked *tHMG* duplicates diverged more from parental genes on autosomes, indicating they experienced different evolutionary forces than the parental copies. Among 243 aligned nucleotide sites, we found 19 nonsynonymous changes and only three synonymous changes shared in all X-linked copies after they diverged from the parental copy (Fig 4B). Similarly, four nonsynonymous changes and no synonymous change occurred on the parental copy in the ancestral species of the *simulans* clade (Fig 4B). The parental copies and new X-linked copies in *D. simulans* and *D. mauritiana* only share ∼70% protein identity, which is very low given the <3 MY divergence. Our branch test using PAML further shows that both branches have significantly higher protein evolution rates (Fig 4C; ω=1.6, LRT test, P= 0.007; Table S10). However, we did not find thedence of positive selection using a branch-site test (LRT test, P= 0.23; Table S10).

### The relocation of ancestrally Y-linked genes from X to Y in the *montium* group species

Although many of these genes are still X-linked in some *montium* group species (*e.g., D. triauraria),* subsequent translocation of some ancestrally Y-linked genes back to the Y chromosome occurred in other species (*e.g., PRY* and *Ppr-Y* in *D. kikkawai*) [12]. Phylogenetic analyses allowed us to unambiguously rule out the alternate hypothesis that following gene duplication onto the X, some lineages lost their Y-linked copies whereas others lost X-linked copies, for the *PRY* and *Ppr-Y* genes (Fig S6). However, we cannot unambiguously distinguish between these two competing hypotheses for the *WDY* and *kl-2* genes in *D. jambulina, D. bocqueti,* and *D. kikkawai* (Fig S6).

## References

1. Raman P, Rominger MC, Young JM, Molaro A, Tsukiyama T, Malik HS. Novel Classes and Evolutionary Turnover of Histone H2B Variants in the Mammalian Germline. Mol Biol Evol. 2022;39(2). Epub 2022/02/01. doi: 10.1093/molbev/msac019. PubMed PMID: 35099534; PubMed Central PMCID: PMCPMC8857922.

2. Molaro A, Wood AJ, Janssens D, Kindelay SM, Eickbush MT, Wu S, et al. Biparental contributions of the H2A.B histone variant control embryonic development in mice. PLoS Biol. 2020;18(12):e3001001. Epub 2020/12/29. doi: 10.1371/journal.pbio.3001001. PubMed PMID: 33362208; PubMed Central PMCID: PMCPMC7757805 following competing interests: HSM is a member of the PLOS Biology editorial board.

3. Talbert PB, Henikoff S. Histone variants at a glance. J Cell Sci. 2021;134(6). Epub 2021/03/28. doi: 10.1242/jcs.244749. PubMed PMID: 33771851; PubMed Central PMCID: PMCPMC8015243.

4. Sassone-Corsi P. Unique chromatin remodeling and transcriptional regulation in spermatogenesis. Science. 2002;296(5576):2176–8. Epub 2002/06/22. doi: 10.1126/science.1070963. PubMed PMID: 12077401.

5. Ward WS, Coffey DS. DNA packaging and organization in mammalian spermatozoa: comparison with somatic cells. Biol Reprod. 1991;44(4):569–74. Epub 1991/04/01. doi: 10.1095/biolreprod44.4.569. PubMed PMID: 2043729.

6. Torok A, Schiffer PH, Schnitzler CE, Ford K, Mullikin JC, Baxevanis AD, et al. The cnidarian Hydractinia echinata employs canonical and highly adapted histones to pack its DNA. Epigenetics Chromatin. 2016;9(1):36. Epub 2016/09/08. doi: 10.1186/s13072-016-0085-1. PubMed PMID: 27602058; PubMed Central PMCID: PMCPMC5011920.

7. Eirin-Lopez JM, Ausio J. Origin and evolution of chromosomal sperm proteins. Bioessays. 2009;31(10):1062–70. Epub 2009/08/27. doi: 10.1002/bies.200900050. PubMed PMID: 19708021.

8. Brewer LR, Corzett M, Balhorn R. Protamine-induced condensation and decondensation of the same DNA molecule. Science. 1999;286(5437):120-3. Epub 1999/10/03. doi: 10.1126/science.286.5437.120. PubMed PMID: 10506559.

9. Luke L, Campbell P, Varea Sanchez M, Nachman MW, Roldan ER. Sexual selection on protamine and transition nuclear protein expression in mouse species. Proc Biol Sci. 2014;281(1783):20133359. Epub 2014/03/29. doi: 10.1098/rspb.2013.3359. PubMed PMID: 24671975; PubMed Central PMCID: PMCPMC3996607.

10. Lupold S, Manier MK, Puniamoorthy N, Schoff C, Starmer WT, Luepold SH, et al. How sexual selection can drive the evolution of costly sperm ornamentation. Nature. 2016;533(7604):535–8. Epub 2016/05/27. doi: 10.1038/nature18005. PubMed PMID: 27225128.

11. Balhorn R. The protamine family of sperm nuclear proteins. Genome Biol. 2007;8(9):227. Epub 2007/10/02. doi: 10.1186/gb-2007-8-9-227. PubMed PMID: 17903313; PubMed Central PMCID: PMCPMC2375014.

12. Martin-Coello J, Gomendio M, Roldan ER. Protamine 3 shows evidence of weak, positive selection in mouse species (genus Mus)--but it is not a protamine. Biol Reprod. 2011;84(2):320–6. Epub 2010/10/15. doi: 10.1095/biolreprod.110.086454. PubMed PMID: 20944085.

13. Nayernia K, Adham I, Kremling H, Reim K, Schlicker M, Schluter G, et al. Stage and developmental specific gene expression during mammalian spermatogenesis. Int J Dev Biol. 1996;40(1):379–83. Epub 1996/02/01. PubMed PMID: 8735951.

14. Cho C, Jung-Ha H, Willis WD, Goulding EH, Stein P, Xu Z, et al. Protamine 2 deficiency leads to sperm DNA damage and embryo death in mice. Biol Reprod. 2003;69(1):211–7. Epub 2003/03/07. doi: 10.1095/biolreprod.102.015115. PubMed PMID: 12620939.

15. Cho C, Willis WD, Goulding EH, Jung-Ha H, Choi YC, Hecht NB, et al. Haploinsufficiency of protamine-1 or -2 causes infertility in mice. Nat Genet. 2001;28(1):82–6. Epub 2001/04/28. doi: 10.1038/ng0501-82. PubMed PMID: 11326282.

16. Reynolds WF, Wolfe SL. Protamines in plant sperm. Exp Cell Res. 1984;152(2):443–8. Epub 1984/06/01. doi: 10.1016/0014-4827(84)90645-1. PubMed PMID: 6723798.

17. Torok A, Gornik SG. Sperm Nuclear Basic Proteins of Marine Invertebrates. Results Probl Cell Differ. 2018;65:15–32. Epub 2018/08/08. doi: 10.1007/978-3-319-92486-1_2. PubMed PMID: 30083913.

18. Lewis JD, Saperas N, Song Y, Zamora MJ, Chiva M, Ausio J. Histone H1 and the origin of protamines. Proc Natl Acad Sci U S A. 2004;101(12):4148–52. Epub 2004/03/17. doi: 10.1073/pnas.0308721101. PubMed PMID: 15024099; PubMed Central PMCID: PMCPMC384709.

19. D’Ippolito RA, Minamino N, Rivera-Casas C, Cheema MS, Bai DL, Kasinsky HE, et al. Protamines from liverwort are produced by post-translational cleavage and C-terminal di-aminopropanelation of several male germ-specific H1 histones. The Journal of biological chemistry. 2019;294(44):16364–73. Epub 2019/09/19. doi: 10.1074/jbc.RA119.010316. PubMed PMID: 31527083; PubMed Central PMCID: PMCPMC6827293.

20. Green GR, Poccia DL. Interaction of sperm histone variants and linker DNA during spermiogenesis in the sea urchin. Biochemistry. 1988;27(2):619–25. Epub 1988/01/26. doi: 10.1021/bi00402a019. PubMed PMID: 3349051.

21. Eirin-Lopez JM, Frehlick LJ, Ausio J. Protamines, in the footsteps of linker histone evolution. The Journal of biological chemistry. 2006;281(1):1–4. Epub 2005/10/26. doi: 10.1074/jbc.R500018200. PubMed PMID: 16243843.

22. Saperas N, Ausio J. Sperm nuclear basic proteins of tunicates and the origin of protamines. Biol Bull. 2013;224(3):127–36. Epub 2013/09/03. doi: 10.1086/BBLv224n3p127. PubMed PMID: 23995738.

23. Wyckoff GJ, Wang W, Wu CI. Rapid evolution of male reproductive genes in the descent of man. Nature. 2000;403(6767):304–9. Epub 2000/02/05. doi: 10.1038/35002070. PubMed PMID: 10659848.

24. Luke L, Tourmente M, Roldan ER. Sexual Selection of Protamine 1 in Mammals. Mol Biol Evol. 2016;33(1):174–84. Epub 2015/10/03. doi: 10.1093/molbev/msv209. PubMed PMID: 26429923.

25. Luke L, Tourmente M, Dopazo H, Serra F, Roldan ER. Selective constraints on protamine 2 in primates and rodents. BMC Evol Biol. 2016;16:21. Epub 2016/01/24. doi: 10.1186/s12862-016-0588-1. PubMed PMID: 26801756; PubMed Central PMCID: PMCPMC4724148.

26. Gartner SM, Rothenbusch S, Buxa MK, Theofel I, Renkawitz R, Rathke C, et al. The HMG-box-containing proteins tHMG-1 and tHMG-2 interact during the histone-to-protamine transition in Drosophila spermatogenesis. Eur J Cell Biol. 2015;94(1):46–59. Epub 2014/12/04. doi: 10.1016/j.ejcb.2014.10.005. PubMed PMID: 25464903.

27. Tirmarche S, Kimura S, Sapey-Triomphe L, Sullivan W, Landmann F, Loppin B. Drosophila protamine-like Mst35Ba and Mst35Bb are required for proper sperm nuclear morphology but are dispensable for male fertility. G3 (Bethesda). 2014;4(11):2241–5. Epub 2014/09/23. doi: 10.1534/g3.114.012724. PubMed PMID: 25236732; PubMed Central PMCID: PMCPMC4232549.

28. Kimura S, Loppin B. The Drosophila chromosomal protein Mst77F is processed to generate an essential component of mature sperm chromatin. Open Biol. 2016;6(11). Epub 2016/11/05. doi: 10.1098/rsob.160207. PubMed PMID: 27810970; PubMed Central PMCID: PMCPMC5133442.

29. Jayaramaiah Raja S, Renkawitz-Pohl R. Replacement by Drosophila melanogaster protamines and Mst77F of histones during chromatin condensation in late spermatids and role of sesame in the removal of these proteins from the male pronucleus. Mol Cell Biol. 2005;25(14):6165–77. Epub 2005/07/01. doi: 10.1128/MCB.25.14.6165-6177.2005. PubMed PMID: 15988027; PubMed Central PMCID: PMCPMC1168805.

30. Eren-Ghiani Z, Rathke C, Theofel I, Renkawitz-Pohl R. Prtl99C Acts Together with Protamines and Safeguards Male Fertility in Drosophila. Cell Rep. 2015;13(11):2327–35. Epub 2015/12/18. doi: 10.1016/j.celrep.2015.11.023. PubMed PMID: 26673329.

31. Yamaki T, Yasuda GK, Wakimoto BT. The Deadbeat Paternal Effect of Uncapped Sperm Telomeres on Cell Cycle Progression and Chromosome Behavior in Drosophila melanogaster. Genetics. 2016;203(2):799–816. Epub 2016/04/01. doi: 10.1534/genetics.115.182436. PubMed PMID: 27029731; PubMed Central PMCID: PMCPMC4896195.

32. Rathke C, Barckmann B, Burkhard S, Jayaramaiah-Raja S, Roote J, Renkawitz-Pohl R. Distinct functions of Mst77F and protamines in nuclear shaping and chromatin condensation during Drosophila spermiogenesis. Eur J Cell Biol. 2010;89(4):326–38. Epub 2010/02/09. doi: 10.1016/j.ejcb.2009.09.001. PubMed PMID: 20138392.

33. Rivard EL, Ludwig AG, Patel PH, Grandchamp A, Arnold SE, Berger A, et al. A putative de novo evolved gene required for spermatid chromatin condensation in Drosophila melanogaster. PLoS Genet. 2021;17(9):e1009787. Epub 2021/09/04. doi: 10.1371/journal.pgen.1009787. PubMed PMID: 34478447; PubMed Central PMCID: PMCPMC8445463.

34. Hempel LU, Rathke C, Raja SJ, Renkawitz-Pohl R. In Drosophila, don juan and don juan like encode proteins of the spermatid nucleus and the flagellum and both are regulated at the transcriptional level by the TAF II80 cannonball while translational repression is achieved by distinct elements. Dev Dyn. 2006;235(4):1053–64. Epub 2006/02/16. doi: 10.1002/dvdy.20698. PubMed PMID: 16477641.

35. Harhangi HR, Sun X, Wang YX, Akhmanova A, Miedema K, Heyting C, et al. RADHA--a new male germ line-specific chromosomal protein of Drosophila. Chromosoma. 1999;108(4):235–42. Epub 1999/08/25. doi: 10.1007/s004120050373. PubMed PMID: 10460411.

36. Vedanayagam J, Lin CJ, Lai EC. Rapid evolutionary dynamics of an expanding family of meiotic drive factors and their hpRNA suppressors. Nat Ecol Evol. 2021;5(12):1613–23. Epub 2021/12/05. doi: 10.1038/s41559-021-01592-z. PubMed PMID: 34862477; PubMed Central PMCID: PMCPMC8665063.

37. Muirhead CA, Presgraves DC. Satellite DNA-mediated diversification of a sex-ratio meiotic drive gene family in Drosophila. Nat Ecol Evol. 2021;5(12):1604–12. Epub 2021/09/08. doi: 10.1038/s41559-021-01543-8. PubMed PMID: 34489561.

38. Courret C, Chang CH, Wei KH, Montchamp-Moreau C, Larracuente AM. Meiotic drive mechanisms: lessons from Drosophila. Proc Biol Sci. 2019;286(1913):20191430. Epub 2019/10/24. doi: 10.1098/rspb.2019.1430. PubMed PMID: 31640520; PubMed Central PMCID: PMCPMC6834043.

39. Gingell LF, McLean JR. A Protamine Knockdown Mimics the Function of Sd in Drosophila melanogaster. G3 (Bethesda). 2020;10(6):2111–5. Epub 2020/04/24. doi: 10.1534/g3.120.401307. PubMed PMID: 32321837; PubMed Central PMCID: PMCPMC7263674.

40. Tao Y, Araripe L, Kingan SB, Ke Y, Xiao H, Hartl DL. A sex-ratio meiotic drive system in Drosophila simulans. II: an X-linked distorter. PLoS Biol. 2007;5(11):e293. Epub 2007/11/09. doi: 10.1371/journal.pbio.0050293. PubMed PMID: 17988173; PubMed Central PMCID: PMCPMC2062476.

41. Faulhaber SH. An abnormal sex ratio in Drosophila simulans. Genetics. 1967;56(1):189–213. Epub 1967/05/01. doi: 10.1093/genetics/56.1.189. PubMed PMID: 6035593; PubMed Central PMCID: PMCPMC1211489.

42. Tao Y, Masly JP, Araripe L, Ke Y, Hartl DL. A sex-ratio meiotic drive system in Drosophila simulans. I: an autosomal suppressor. PLoS Biol. 2007;5(11):e292. Epub 2007/11/09. doi: 10.1371/journal.pbio.0050292. PubMed PMID: 17988172; PubMed Central PMCID: PMCPMC2062475.

43. Rathke C, Baarends WM, Jayaramaiah-Raja S, Bartkuhn M, Renkawitz R, Renkawitz-Pohl R. Transition from a nucleosome-based to a protamine-based chromatin configuration during spermiogenesis in Drosophila. J Cell Sci. 2007;120(Pt 9):1689–700. Epub 2007/04/25. doi: 10.1242/jcs.004663. PubMed PMID: 17452629.

44. Suvorov A, Kim BY, Wang J, Armstrong EE, Peede D, D’Agostino ERR, et al. Widespread introgression across a phylogeny of 155 Drosophila genomes. Curr Biol. 2022;32(1):111–23 e5. Epub 2021/11/18. doi: 10.1016/j.cub.2021.10.052. PubMed PMID: 34788634; PubMed Central PMCID: PMCPMC8752469.

45. Weisman CM, Murray AW, Eddy SR. Many, but not all, lineage-specific genes can be explained by homology detection failure. PLoS Biol. 2020;18(11):e3000862. Epub 2020/11/03. doi: 10.1371/journal.pbio.3000862. PubMed PMID: 33137085; PubMed Central PMCID: PMCPMC7660931.

46. Reddiex AJ, Allen SL, Chenoweth SF. A Genomic Reference Panel for Drosophila serrata. G3 (Bethesda). 2018;8(4):1335–46. Epub 2018/03/01. doi: 10.1534/g3.117.300487. PubMed PMID: 29487184; PubMed Central PMCID: PMCPMC5873922.

47. Bronski MJ, Martinez CC, Weld HA, Eisen MB. Whole Genome Sequences of 23 Species from the Drosophila montium Species Group (Diptera: Drosophilidae): A Resource for Testing Evolutionary Hypotheses. G3 (Bethesda). 2020;10(5):1443–55. Epub 2020/03/30. doi: 10.1534/g3.119.400959. PubMed PMID: 32220952; PubMed Central PMCID: PMCPMC7202002.

48. Yang Z. PAML 4: phylogenetic analysis by maximum likelihood. Mol Biol Evol. 2007;24(8):1586–91. Epub 2007/05/08. doi: 10.1093/molbev/msm088. PubMed PMID: 17483113.

49. Yang Z, Nielsen R, Goldman N, Pedersen AM. Codon-substitution models for heterogeneous selection pressure at amino acid sites. Genetics. 2000;155(1):431–49. Epub 2000/05/03. doi: 10.1093/genetics/155.1.431. PubMed PMID: 10790415; PubMed Central PMCID: PMCPMC1461088.

50. Kasinathan B, Colmenares SU, 3rd, McConnell H, Young JM, Karpen GH, Malik HS. Innovation of heterochromatin functions drives rapid evolution of essential ZAD-ZNF genes in Drosophila. Elife. 2020;9. Epub 2020/11/11. doi: 10.7554/eLife.63368. PubMed PMID: 33169670; PubMed Central PMCID: PMCPMC7655104.

51. Bayes JJ, Malik HS. Altered heterochromatin binding by a hybrid sterility protein in Drosophila sibling species. Science. 2009;326(5959):1538–41. Epub 2009/11/26. doi: 10.1126/science.1181756. PubMed PMID: 19933102; PubMed Central PMCID: PMCPMC2987944.

52. Kim BY, Wang JR, Miller DE, Barmina O, Delaney E, Thompson A, et al. Highly contiguous assemblies of 101 drosophilid genomes. Elife. 2021;10. Epub 2021/07/20. doi: 10.7554/eLife.66405. PubMed PMID: 34279216; PubMed Central PMCID: PMCPMC8337076.

53. Altschul SF, Gish W, Miller W, Myers EW, Lipman DJ. Basic local alignment search tool. J Mol Biol. 1990;215(3):403–10. Epub 1990/10/05. doi: 10.1016/S0022-2836(05)80360-2. PubMed PMID: 2231712.

54. Manni M, Berkeley MR, Seppey M, Simao FA, Zdobnov EM. BUSCO Update: Novel and Streamlined Workflows along with Broader and Deeper Phylogenetic Coverage for Scoring of Eukaryotic, Prokaryotic, and Viral Genomes. Mol Biol Evol. 2021;38(10):4647–54. Epub 2021/07/29. doi: 10.1093/molbev/msab199. PubMed PMID: 34320186; PubMed Central PMCID: PMCPMC8476166.

55. Dupim EG, Goldstein G, Vanderlinde T, Vaz SC, Krsticevic F, Bastos A, et al. An investigation of Y chromosome incorporations in 400 species of Drosophila and related genera. PLoS Genet. 2018;14(11):e1007770. Epub 2018/11/06. doi: 10.1371/journal.pgen.1007770. PubMed PMID: 30388103; PubMed Central PMCID: PMCPMC6235401.

56. Conner WR, Delaney EK, Bronski MJ, Ginsberg PS, Wheeler TB, Richardson KM, et al. A phylogeny for the Drosophila montium species group: A model clade for comparative analyses. Mol Phylogenet Evol. 2021;158:107061. Epub 2021/01/03. doi: 10.1016/j.ympev.2020.107061. PubMed PMID: 33387647; PubMed Central PMCID: PMCPMC7946709.

57. Barckmann B, Chen X, Kaiser S, Jayaramaiah-Raja S, Rathke C, Dottermusch-Heidel C, et al. Three levels of regulation lead to protamine and Mst77F expression in Drosophila. Dev Biol. 2013;377(1):33–45. Epub 2013/03/08. doi: 10.1016/j.ydbio.2013.02.018. PubMed PMID: 23466740; PubMed Central PMCID: PMCPMC4154633.

58. Hecht NB, Bower PA, Waters SH, Yelick PC, Distel RJ. Evidence for haploid expression of mouse testicular genes. Exp Cell Res. 1986;164(1):183–90. Epub 1986/05/01. doi: 10.1016/0014-4827(86)90465-9. PubMed PMID: 3754219.

59. Peschon JJ, Behringer RR, Brinster RL, Palmiter RD. Spermatid-specific expression of protamine 1 in transgenic mice. Proc Natl Acad Sci U S A. 1987;84(15):5316–9. Epub 1987/08/01. doi: 10.1073/pnas.84.15.5316. PubMed PMID: 3037541; PubMed Central PMCID: PMCPMC298846.

60. Perreault SD. Chromatin remodeling in mammalian zygotes. Mutat Res. 1992;296(1-2):43–55. Epub 1992/12/01. doi: 10.1016/0165-1110(92)90031-4. PubMed PMID: 1279407.

61. Loppin B, Docquier M, Bonneton F, Couble P. The maternal effect mutation sesame affects the formation of the male pronucleus in Drosophila melanogaster. Dev Biol. 2000;222(2):392–404. Epub 2000/06/06. doi: 10.1006/dbio.2000.9718. PubMed PMID: 10837127.

62. Beckmann JF, Sharma GD, Mendez L, Chen H, Hochstrasser M. The Wolbachia cytoplasmic incompatibility enzyme CidB targets nuclear import and protamine-histone exchange factors. Elife. 2019;8. Epub 2019/11/28. doi: 10.7554/eLife.50026. PubMed PMID: 31774393; PubMed Central PMCID: PMCPMC6881146.

63. Kaur R, Leigh BA, Ritchie IT, Bordenstein SR. The Cif proteins from Wolbachia prophage WO modify sperm genome integrity to establish cytoplasmic incompatibility. PLoS Biol. 2022;20(5):e3001584. Epub 2022/05/25. doi: 10.1371/journal.pbio.3001584. PubMed PMID: 35609042.

64. Horard B, Terretaz K, Gosselin-Grenet AS, Sobry H, Sicard M, Landmann F, et al. Paternal transmission of the Wolbachia CidB toxin underlies cytoplasmic incompatibility. Curr Biol. 2022;32(6):1319–31 e5. Epub 2022/02/09. doi: 10.1016/j.cub.2022.01.052. PubMed PMID: 35134330.

65. Sturtevant AH, Novitski E. The Homologies of the Chromosome Elements in the Genus Drosophila. Genetics. 1941;26(5):517–41. Epub 1941/09/01. doi: 10.1093/genetics/26.5.517. PubMed PMID: 17247021; PubMed Central PMCID: PMCPMC1209144.

66. Chang T-P, Tsai T-H, Chang H-y. Fusions of Muller’s Elements during Chromosome Evolution of Drosophila albomicans. Zoological Studies. 2010;49(1):152.

67. Zhou Q, Zhu HM, Huang QF, Zhao L, Zhang GJ, Roy SW, et al. Deciphering neo-sex and B chromosome evolution by the draft genome of Drosophila albomicans. BMC Genomics. 2012;13:109. Epub 2012/03/24. doi: 10.1186/1471-2164-13-109. PubMed PMID: 22439699; PubMed Central PMCID: PMCPMC3353239.

68. Ellison C, Bachtrog D. Recurrent gene co-amplification on Drosophila X and Y chromosomes. PLoS Genet. 2019;15(7):e1008251. Epub 2019/07/23. doi: 10.1371/journal.pgen.1008251. PubMed PMID: 31329593; PubMed Central PMCID: PMCPMC6690552.

69. Saint-Leandre B, Levine MT. The Telomere Paradox: Stable Genome Preservation with Rapidly Evolving Proteins. Trends Genet. 2020;36(4):232–42. Epub 2020/03/11. doi: 10.1016/j.tig.2020.01.007. PubMed PMID: 32155445; PubMed Central PMCID: PMCPMC7066039.

70. Levine MT, Vander Wende HM, Malik HS. Mitotic fidelity requires transgenerational action of a testis-restricted HP1. Elife. 2015;4:e07378. Epub 2015/07/08. doi: 10.7554/eLife.07378. PubMed PMID: 26151671; PubMed Central PMCID: PMCPMC4491702.

71. Ross BD, Rosin L, Thomae AW, Hiatt MA, Vermaak D, de la Cruz AF, et al. Stepwise evolution of essential centromere function in a Drosophila neogene. Science. 2013;340(6137):1211–4. Epub 2013/06/08. doi: 10.1126/science.1234393. PubMed PMID: 23744945; PubMed Central PMCID: PMCPMC4119826.

72. Drosophila 12 Genomes C, Clark AG, Eisen MB, Smith DR, Bergman CM, Oliver B, et al. Evolution of genes and genomes on the Drosophila phylogeny. Nature. 2007;450(7167):203–18. Epub 2007/11/13. doi: 10.1038/nature06341. PubMed PMID: 17994087.

73. Minh BQ, Schmidt HA, Chernomor O, Schrempf D, Woodhams MD, von Haeseler A, et al. IQ-TREE 2: New Models and Efficient Methods for Phylogenetic Inference in the Genomic Era. Mol Biol Evol. 2020;37(5):1530–4. Epub 2020/02/06. doi: 10.1093/molbev/msaa015. PubMed PMID: 32011700; PubMed Central PMCID: PMCPMC7182206.

74. McDonald JH, Kreitman M. Adaptive protein evolution at the Adh locus in Drosophila. Nature. 1991;351(6328):652–4. Epub 1991/06/20. doi: 10.1038/351652a0. PubMed PMID: 1904993.

75. Stecher G, Tamura K, Kumar S. Molecular Evolutionary Genetics Analysis (MEGA) for macOS. Mol Biol Evol. 2020;37(4):1237–9. Epub 2020/01/07. doi: 10.1093/molbev/msz312. PubMed PMID: 31904846; PubMed Central PMCID: PMCPMC7086165.

76. Hervas S, Sanz E, Casillas S, Pool JE, Barbadilla A. PopFly: the Drosophila population genomics browser. Bioinformatics. 2017;33(17):2779–80. Epub 2017/05/05. doi: 10.1093/bioinformatics/btx301. PubMed PMID: 28472360; PubMed Central PMCID: PMCPMC5860067.

77. Lack JB, Lange JD, Tang AD, Corbett-Detig RB, Pool JE. A Thousand Fly Genomes: An Expanded Drosophila Genome Nexus. Mol Biol Evol. 2016;33(12):3308–13. Epub 2016/10/01. doi: 10.1093/molbev/msw195. PubMed PMID: 27687565; PubMed Central PMCID: PMCPMC5100052.

78. Egea R, Casillas S, Barbadilla A. Standard and generalized McDonald-Kreitman test: a website to detect selection by comparing different classes of DNA sites. Nucleic Acids Res. 2008;36(Web Server issue):W157–62. Epub 2008/06/03. doi: 10.1093/nar/gkn337. PubMed PMID: 18515345; PubMed Central PMCID: PMCPMC2447769.

79. Kim D, Paggi JM, Park C, Bennett C, Salzberg SL. Graph-based genome alignment and genotyping with HISAT2 and HISAT-genotype. Nat Biotechnol. 2019;37(8):907–15. Epub 2019/08/04. doi: 10.1038/s41587-019-0201-4. PubMed PMID: 31375807; PubMed Central PMCID: PMCPMC7605509.

80. Kovaka S, Zimin AV, Pertea GM, Razaghi R, Salzberg SL, Pertea M. Transcriptome assembly from long-read RNA-seq alignments with StringTie2. Genome Biol. 2019;20(1):278. Epub 2019/12/18. doi: 10.1186/s13059-019-1910-1. PubMed PMID: 31842956; PubMed Central PMCID: PMCPMC6912988.

81. Li H. New strategies to improve minimap2 alignment accuracy. Bioinformatics. 2021. Epub 2021/10/09. doi: 10.1093/bioinformatics/btab705. PubMed PMID: 34623391; PubMed Central PMCID: PMCPMC8652018.

82. Vasimuddin M, Misra S, Li H, Aluru S, editors. Efficient Architecture-Aware Acceleration of BWA-MEM for Multicore Systems. 2019. IEEE International Parallel and Distributed Processing Symposium (IPDPS); 2019 20–24 May 2019.

83. Chang CH, Larracuente AM. Heterochromatin-Enriched Assemblies Reveal the Sequence and Organization of the Drosophila melanogaster Y Chromosome. Genetics. 2019;211(1):333–48. Epub 2018/11/14. doi: 10.1534/genetics.118.301765. PubMed PMID: 30420487; PubMed Central PMCID: PMCPMC6325706.

84. Li H, Handsaker B, Wysoker A, Fennell T, Ruan J, Homer N, et al. The Sequence Alignment/Map format and SAMtools. Bioinformatics. 2009;25(16):2078–9. Epub 2009/06/10. doi: 10.1093/bioinformatics/btp352. PubMed PMID: 19505943; PubMed Central PMCID: PMCPMC2723002.

85. Li H. A statistical framework for SNP calling, mutation discovery, association mapping and population genetical parameter estimation from sequencing data. Bioinformatics. 2011;27(21):2987–93. Epub 2011/09/10. doi: 10.1093/bioinformatics/btr509. PubMed PMID: 21903627; PubMed Central PMCID: PMCPMC3198575.

86. Bellen HJ, Levis RW, Liao G, He Y, Carlson JW, Tsang G, et al. The BDGP gene disruption project: single transposon insertions associated with 40% of Drosophila genes. Genetics. 2004;167(2):761–81. Epub 2004/07/09. doi: 10.1534/genetics.104.026427. PubMed PMID: 15238527; PubMed Central PMCID: PMCPMC1470905.

87. Parks AL, Cook KR, Belvin M, Dompe NA, Fawcett R, Huppert K, et al. Systematic generation of high-resolution deletion coverage of the Drosophila melanogaster genome. Nat Genet. 2004;36(3):288–92. Epub 2004/02/26. doi: 10.1038/ng1312. PubMed PMID: 14981519.

88. Kumar S, Stecher G, Suleski M, Hedges SB. TimeTree: A Resource for Timelines, Timetrees, and Divergence Times. Mol Biol Evol. 2017;34(7):1812–9. Epub 2017/04/08. doi: 10.1093/molbev/msx116. PubMed PMID: 28387841.

89. Krsticevic FJ, Schrago CG, Carvalho AB. Long-Read Single Molecule Sequencing to Resolve Tandem Gene Copies: The Mst77Y Region on the Drosophila melanogaster Y Chromosome. G3 (Bethesda). 2015;5(6):1145–50. Epub 2015/04/11. doi: 10.1534/g3.115.017277. PubMed PMID: 25858959; PubMed Central PMCID: PMCPMC4478544.

90. Chakraborty M, Chang CH, Khost DE, Vedanayagam J, Adrion JR, Liao Y, et al. Evolution of genome structure in the Drosophila simulans species complex. Genome Res. 2021;31(3):380–96. Epub 2021/02/11. doi: 10.1101/gr.263442.120. PubMed PMID: 33563718; PubMed Central PMCID: PMCPMC7919458.

## References

1. Hervas S, Sanz E, Casillas S, Pool JE, Barbadilla A. PopFly: the Drosophila population genomics browser. Bioinformatics. 2017;33(17):2779–80. Epub 2017/05/05. doi: 10.1093/bioinformatics/btx301. PubMed PMID: 28472360; PubMed Central PMCID: PMCPMC5860067.

2. Mackay TF, Richards S, Stone EA, Barbadilla A, Ayroles JF, Zhu D, et al. The Drosophila melanogaster Genetic Reference Panel. Nature. 2012;482(7384):173–8. Epub 2012/02/10. doi: 10.1038/nature10811. PubMed PMID: 22318601; PubMed Central PMCID: PMCPMC3683990.

3. Campo D, Lehmann K, Fjeldsted C, Souaiaia T, Kao J, Nuzhdin SV. Whole-genome sequencing of two North American Drosophila melanogaster populations reveals genetic differentiation and positive selection. Mol Ecol. 2013;22(20):5084–97. Epub 2013/10/10. doi: 10.1111/mec.12468. PubMed PMID: 24102956; PubMed Central PMCID: PMCPMC3800041.

4. Kao JY, Zubair A, Salomon MP, Nuzhdin SV, Campo D. Population genomic analysis uncovers African and European admixture in Drosophila melanogaster populations from the south-eastern United States and Caribbean Islands. Mol Ecol. 2015;24(7):1499–509. Epub 2015/03/05. doi: 10.1111/mec.13137. PubMed PMID: 25735402.

5. Grenier JK, Arguello JR, Moreira MC, Gottipati S, Mohammed J, Hackett SR, et al. Global diversity lines – a five-continent reference panel of sequenced Drosophila melanogaster strains. G3 (Bethesda). 2015;5(4):593–603. Epub 2015/02/13. doi: 10.1534/g3.114.015883. PubMed PMID: 25673134; PubMed Central PMCID: PMCPMC4390575.

6. Lack JB, Lange JD, Tang AD, Corbett-Detig RB, Pool JE. A Thousand Fly Genomes: An Expanded Drosophila Genome Nexus. Mol Biol Evol. 2016;33(12):3308–13. Epub 2016/10/01. doi: 10.1093/molbev/msw195. PubMed PMID: 27687565; PubMed Central PMCID: PMCPMC5100052.

7. King EG, Macdonald SJ, Long AD. Properties and power of the Drosophila Synthetic Population Resource for the routine dissection of complex traits. Genetics. 2012;191(3):935–49. Epub 2012/04/17. doi: 10.1534/genetics.112.138537. PubMed PMID: 22505626; PubMed Central PMCID: PMCPMC3389985.

8. Langley CH, Stevens K, Cardeno C, Lee YC, Schrider DR, Pool JE, et al. Genomic variation in natural populations of Drosophila melanogaster. Genetics. 2012;192(2):533–98. Epub 2012/06/08. doi: 10.1534/genetics.112.142018. PubMed PMID: 22673804; PubMed Central PMCID: PMCPMC3454882.

9. Bergman CM, Haddrill PR. Strain-specific and pooled genome sequences for populations of Drosophila melanogaster from three continents. F1000Res. 2015;4:31. Epub 2015/02/27. doi: 10.12688/f1000research.6090.1. PubMed PMID: 25717372; PubMed Central PMCID: PMCPMC4331666.

10. Huang W, Massouras A, Inoue Y, Peiffer J, Ramia M, Tarone AM, et al. Natural variation in genome architecture among 205 Drosophila melanogaster Genetic Reference Panel lines. Genome Res. 2014;24(7):1193–208. Epub 2014/04/10. doi: 10.1101/gr.171546.113. PubMed PMID: 24714809; PubMed Central PMCID: PMCPMC4079974.

11. Hurst LD, Smith NGC. The evolution of concerted evolution. Proceedings of the Royal Society of London Series B: Biological Sciences. 1998;265(1391):121–7. doi: 10.1098/rspb.1998.0272.

12. Dupim EG, Goldstein G, Vanderlinde T, Vaz SC, Krsticevic F, Bastos A, et al. An investigation of Y chromosome incorporations in 400 species of Drosophila and related genera. PLoS Genet. 2018;14(11):e1007770. Epub 2018/11/06. doi: 10.1371/journal.pgen.1007770. PubMed PMID: 30388103; PubMed Central PMCID: PMCPMC6235401.

